# Evidence that interspecies *Leishmania* hybrids contribute to changes in disease pathology

**DOI:** 10.1101/2020.06.29.177667

**Authors:** Patrick Lypaczewski, Greg Matlashewski

## Abstract

**Background:** Leishmaniasis is a widespread neglected tropical disease present in over 90 countries with diverse pathologies associated with different species of *Leishmania* parasites transmitted by infected sand flies. *Leishmania donovani* causes visceral leishmaniasis, a highly virulent fatal infection of the visceral organs. *Leishmania major* and *Leishmania tropica* cause less virulent cutaneous leishmaniasis where the infection remains in the skin at the site of the sandfly bite. A major molecular epidemiological question is why some variants of *L. donovani* in Sri Lanka cause cutaneous disease rather than the typical visceral disease.

**Methods:** Whole genome sequencing data for 684 *L. donovani* samples was used to perform sequence alignments and worldwide phylogenetic analyses to determine the source of the atypical *L. donovani* strains from Sri Lanka. *L. donovani* genome sequences originating from Sri Lanka were further analyzed for evidence of hybridization with other *Leishmania* species by determining the density of heterozygous alleles. Polymorphisms from potential *Leishmania* hybrids were used to reconstruct the parental genetic sequences to identify the potential parental species and quantify their genetic contribution through sequence comparison of the reconstructed parental sequences with all Old World *Leishmania* genomes.

**Findings:** Here we show that *L. donovani* in Sri Lanka contains genes with widespread gene polymorphisms derived from African *L. major* and *L. tropica* genomes that were likely obtained as a result of diploid genome hybridization and recombination resulting in progeny with mosaic genomes. Furthermore, evidence is presented that multiple *L. donovani* hybrid parasites originating from visceral leishmaniasis endemic Africa have entered Sri Lanka yet visceral leishmaniasis remains non-existent raising the possibility that environmental factors favour the establishment of atypical *L. donovani* strains in Sri Lanka.

**Interpretation:** The discovery of *L. major* and *L. tropica* genome sequences in *L. donovani* provides a compelling rationale how some *L. donovani* strains in Sri Lanka may be able to cause cutaneous rather than visceral leishmaniasis. The identification of *L. donovani* hybrid parasites in cutaneous leishmaniasis lesions provides a unique opportunity to investigate environmental and parasite genetic factors controlling disease epidemiology and pathogenesis.

**Funding:** Canadian Institutes of Health Research and Fonds de recherche du Québec – Santé

**Research in context:** *Evidence before this study:* Different *Leishmania* species parasites cause either benign cutaneous leishmaniasis or fatal visceral leishmaniasis. It is unknown why some variants of *Leishmania donovani* that typically causes visceral leishmaniasis in Asia and Africa can cause cutaneous leishmaniasis in specific geographic locations including Sri Lanka. *Leishmania* has a diploid genome and hybrid parasites have been identified in nature and generated experimentally. In the context of this study, hybrids are considered to be progeny derived from a single outcross event between two diverse parents. Uncertainty remains whether interspecies hybrids with visceral and cutaneous leishmaniasis causing species in nature are associated with different disease outcomes.

*Added value of this study:* Evidence for genetic hybridization between visceral and cutaneous disease causing *Leishmania* species is described from Sri Lanka where cutaneous leishmaniasis is highly endemic yet there is no ongoing visceral leishmaniasis transmission. This provides a potential explanation how *L. donovani* can become attenuated for visceral disease and could help to identify geographic environmental factors associated with selection for parasite attenuation.

*Implications of all the available evidence:* Hybrid *Leishmania* parasites may be one source of atypical cutaneous leishmaniasis. Epidemiological studies are needed to determine why diverse *L. donovani* hybrid parasites have become ubiquitous in specific geographic locations where the incidence of cutaneous leishmaniasis is increasing. This has implications for understanding the genetic control of disease pathogenesis and for the prevention of cutaneous or visceral leishmaniasis locally and in neighboring countries.

## Introduction

Leishmaniasis is a neglected tropical disease present throughout the tropics and subtropics and is caused by the protozoan parasite from the genus *Leishmania* that is transmitted by infected sand flies^1,2^. There are two major pathologic forms associated with leishmaniasis; cutaneous leishmaniasis results in skin lesions at the site of the sand fly bite that usually self-heal within several weeks or months, and the more virulent form, visceral leishmaniasis where *Leishmania* infects the visceral organs and is fatal if not treated. Visceral leishmaniasis is the second most deadly vector borne parasitic diseases after malaria^2,3^. The highest incidence of visceral leishmaniasis is in South East Asia and Sub-Saharan Africa where the predominant etiologic agent is *L. donovani* transmitted by *Phlebotomus spp* sand flies and humans are the only known reservoir^2,3^. Cutaneous leishmaniasis is more widespread internationally than visceral leishmaniasis and is caused by numerous *Leishmania* species, all of which have animal reservoirs with the exception of *L. tropica* that has mainly human reservoirs with some animal reservoirs^2,4^.

Hybrids between related parasites from the *L. donovani* complex strains have been described^5–9^ and there is evidence for hybrids between distant species such as *L. infantum* and *L. major* among natural isolates^10–13^. In the context of this study, hybrids are considered to be progeny derived from a single outcross event between two diverse parents. Experimentally, inter- and intraspecies hybrids have been generated in the sand fly vector, confirming that the promastigote stage of *Leishmania* can form hybrids and carry out genetic exchange^14–17^. More recently, classical chromosome crossing over during meiotic-like recombination and the ability to experimentally conduct backcrosses with F1 progeny has been demonstrated with intraspecies hybrids in sand flies^18^. These observations confirm that intra and inter-species genetic exchange can occur both experimentally and in nature. Uncertainty remains however to what extent interspecies hybrids can contribute to the rise of parasites with different epidemiology and pathogenesis in nature, such as for example *L. donovani* and *L. major* hybrids associated with cutaneous leishmaniasis.

Cutaneous leishmaniasis has recently become endemic in Sri Lanka and there have been over 15,000 cases since 2001 with over 3000 reported cases in 2018^19,20^. In contrast, there have been only 7 suspected cases of visceral leishmaniasis in Sri Lanka since 2004 mostly in individuals with co-morbidities associated with immunosuppression^21^. No confirmed cases of visceral leishmaniasis from Sri Lanka have ever been reported to the WHO^22^. Cutaneous leishmaniasis in Sri Lanka is unique and interesting because it is caused by *L. donovani* which typically causes visceral leishmaniasis in other countries, although recently cutaneous leishmaniasis caused by *L. donovani* has also been observed in some regions of India and Nepal ^20,23^.

Ongoing research in our laboratory has identified a number of non-synonymous single nucleotide polymorphisms (SNPs) and copy number variations in a Sri Lankan *L. donovani* isolate causing cutaneous leishmaniasis^24,25^. These genetic changes have resulted in biological changes in the conserved mTOR signalling pathway and a reduction in A2 virulence genes that in addition to other unidentified mechanisms, contribute to the atypical cutaneous leishmaniasis phenotype of *L. donovani* in Sri Lanka ^24,25^. More recently, genome sequences from additional *L. donovani* isolates from Sri Lanka have been reported and deposited in NCBI GenBank^26^. Interestingly, these newer isolates did not contain any of the sequence variations and SNPs that we previously identified in the cutaneous leishmaniasis isolate^24,25^ suggesting the presence of multiple co-existing *L. donovani* strains in Sri Lanka. It was therefore necessary to investigate the origin of the *L. donovani* strains in Sri Lanka to understand the etiologic origin of cutaneous leishmaniasis on this island.

Results from this study demonstrate that the *L. donovani* genomes from some strains present in Sri Lanka were remarkably divergent, and evidence is presented here that this is largely due to the presence of parasites with hybrid- genomes including *L. donovani/L. major* and *L. donovani/L. tropica* hybrids. These findings provide one possible explanation for the atypical cutaneous leishmaniasis phenotype in Sri Lanka.

## Results

### Global distribution of *L. donovani* sequences and the divergence of strains from Sri Lanka

To understand the geographic origins of the *L. donovani* strains circulating in Sri Lanka, we compared their genomes to all available *L. donovani* genomes in GenBank^27^ including relevant recent contributions^28^. The entirety of the NCBI GenBank/Sequencing Read Archive (SRA) records for *L. donovani* were data mined as detailed in Methods. As shown in Figure 1A, only whole genome sequencing projects with sufficient quality were included to generate variant profiles for comparative genomics and phylogeny analysis. From the resulting 684 filtered genome sequences, a neighbor-joining phylogenetic tree was generated as shown in Figure 1B. The tree generated in Figure 1B is also available in an interactive format with branches identified by their GenBank SRA accession codes at https://itol.embl.de/tree/1322162673368791580134755, and the accession codes also listed in Supplementary Table S1. The available sequences from the Sri Lanka *L. donovani* isolates formed three distinct groups termed SL1, SL2 and SL3. The SL1 group was closer to the Indian subcontinent group and was comprised of strains originally isolated from Sri Lanka almost 10 years ago^24,25^, as well as sequences from an independent group (NCBI BioProject PRJEB2600). Five of the Sri Lankan *L. donovani* genomes clustered in SL2 were much further than any other *L. donovani* cluster from the Indian subcontinent or from Africa. Three genomes clustered in the SL3 group and were on the edge of the Sudanese/North-Ethiopian *L. donovani* cluster. This demonstrates that the Sri Lanka isolates in groups SL2 and SL3, despite being geographically close to India, are quite different in origin from the SL1 group and that the SL2 group is very unique from any other *L. donovani* strain. All Indian subcontinent (ISC) genomes cluster closely together including the slightly divergent ISC1 or “Yeti” group consistent with previous phylogenetic analyses^29,30^. Parasites from Africa formed three separate clusters, largely determined by their geographical isolation location, as previously reported^7^. Parasites from the north of Ethiopia and Sudan formed a cluster distinct from southern Ethiopian and Kenyan clusters with hybrid parasites between the north and south of Ethiopia forming a small intermediate cluster (highlighted in blue)^6–9,31^.

**Figure 1.**
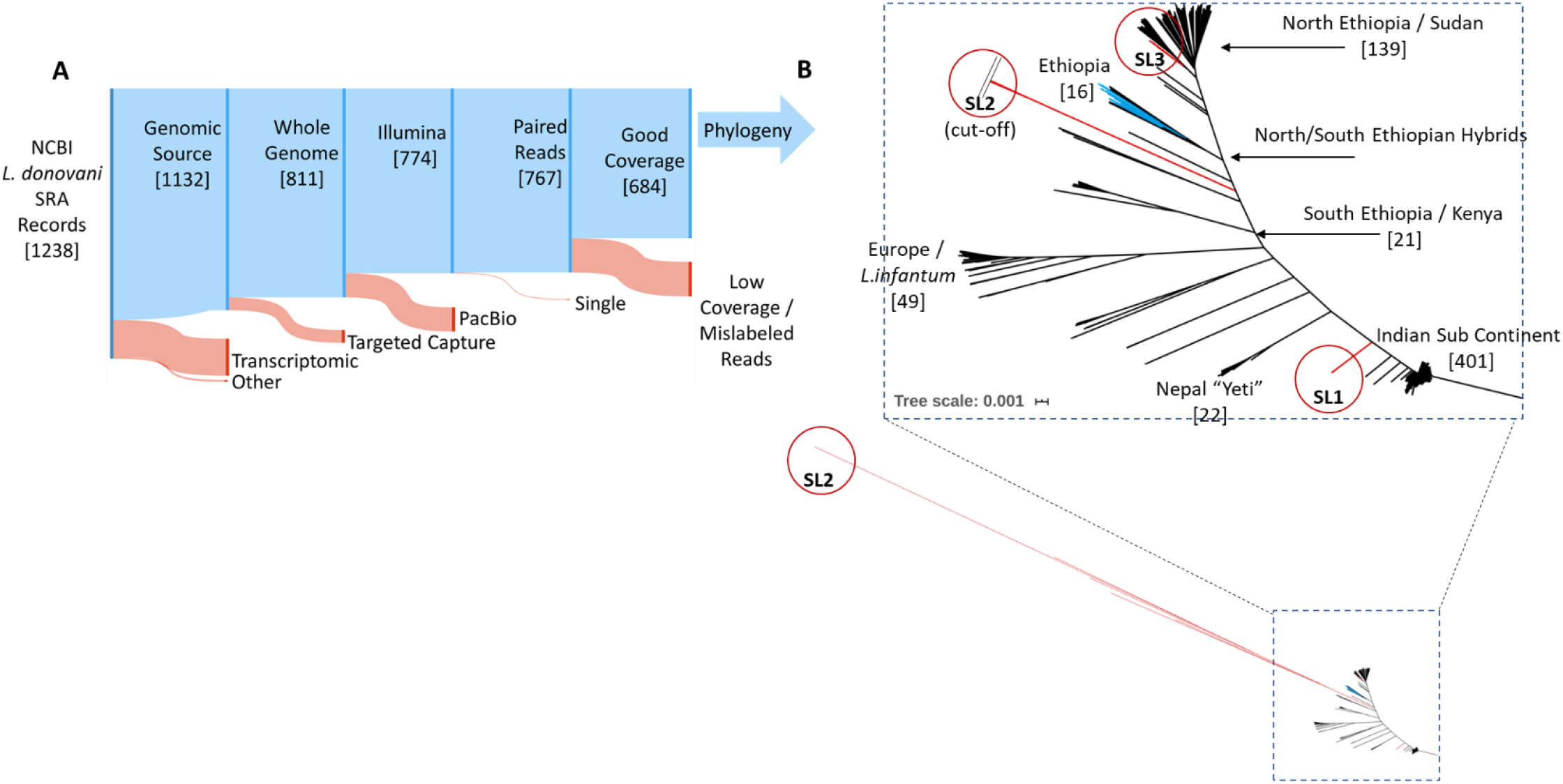
Worldwide *L. donovani* phylogenetic tree. **A.** Selection filters applied to the NCBI 1242 public *L. donovani* Sequence Read Archive (SRA) records to obtain good quality *L. donovani* sequencing data for phylogenetic analysis. Sequencing records were retained if they originated from genomic DNA, had no selection bias for genomic location, were sequenced on high accuracy Illumina platforms in paired sequencing mode and had coverage spanning the entire genome. **B.** Neighbor Joining based tree of all *L. donovani* samples analyzed showing clear geographical groupings. 401 Indian Sub Continent samples cluster close to each other and next to the previously sequenced Sri Lankan isolates^26^ labeled **SL1** (Sri Lanka Group 1) and the Nepalese highland “Yeti” strains. Ethiopian isolates form three separate clusters based on genotype, North, South and hybrid (blue). Five highly divergent Sri Lankan Sri Lanka Group 2 isolates (**SL2**) cluster at a long distance from all other *L. donovani* isolates. Three Sri Lanka Group 3 (**SL3**) isolates cluster on the edge of the North Ethiopian cluster.

As one of the Sri Lankan groups (SL2) diverged more than the African to Indian genetic distances based on branch length (Fig. 1A), we manually inspected the alignments to investigate how this may have occurred. It became apparent that the genomes from the SL2 group were heavily populated with SNPs occurring in the 40-60% frequency range and this was not the case for the SL1 and SL3 groups. For clarity, a representative example of these types of SNPs is shown for a 40 bp section of chromosome 1 (Fig. 2A). The frequency of SNPs is also shown for the entire chromosome 1 (Supplementary Figure S1A) and across the entire genome (Supplementary Figure S1B, S1C, Supplementary Table S2). These data show the different levels of heterozygosity across the entire genome for groups SL1, SL2 and SL3 with group SL2 in the 50% frequency range for diploid chromosomes while group SL3 genomes contain mainly homozygous polymorphisms located on the outer edge of each track. The high frequency heterozygosity in the 50% range was atypical and could represent regions with equal contributions from homologous chromosomes from different parasite genomes and could explain why the SL2 group is phylogenetically very different from other *L. donovani* strains.

**Figure 2.**
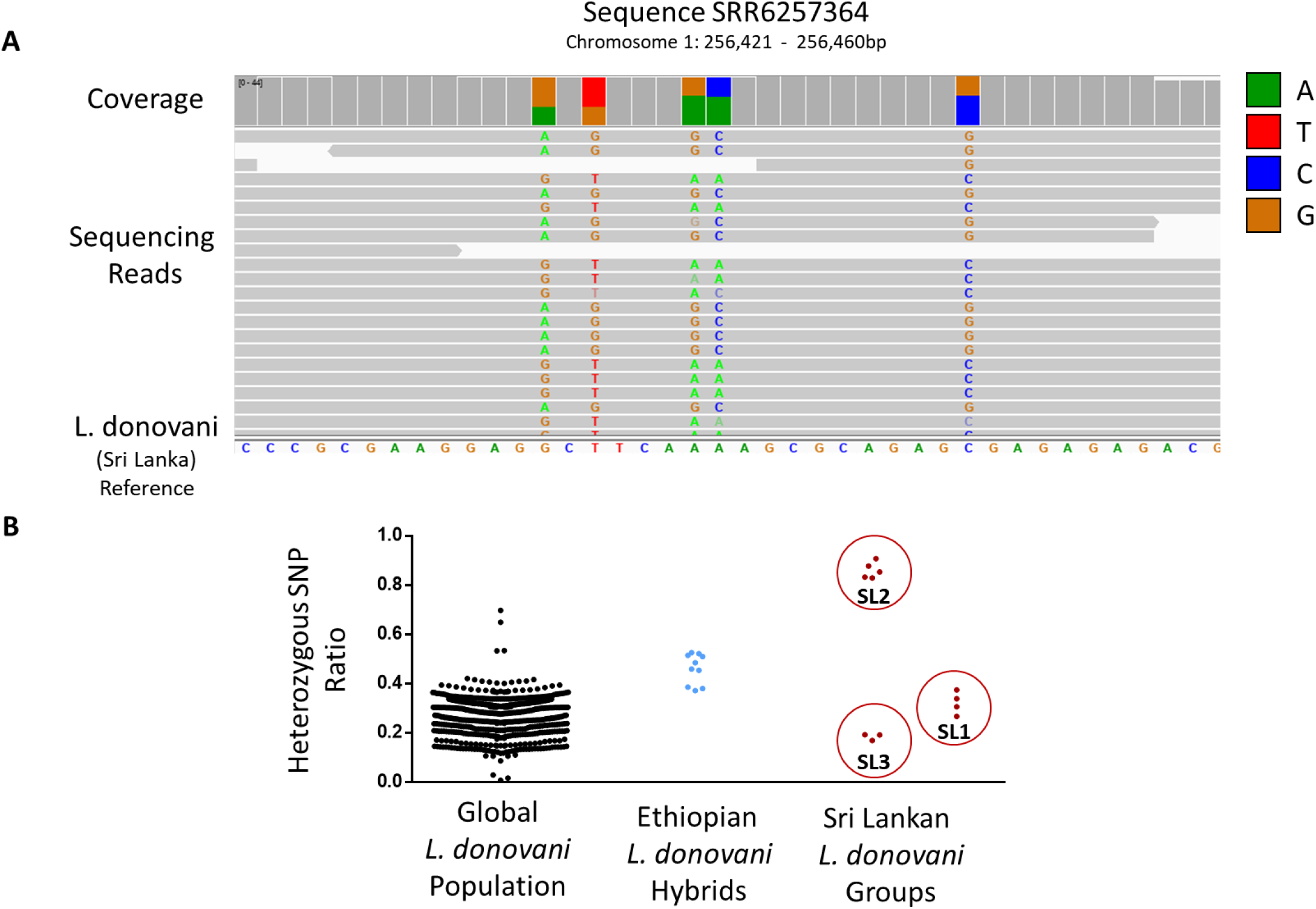
Distribution of heterozygous polymorphisms across all samples. **A.** Representative alignment of SL2 group samples on a 40bp region of *L. donovani* chromosome 1 showing frequent heterozygosity. Reads matching the reference sequence^24^ (bottom) with no SNPs are displayed in gray. Loci where reads contain variability are highlighted in colors corresponding to the respective nucleotides. **B.** Comparison of the heterozygous SNP frequency in all *L. donovani* world wide isolates, known Ethiopian hybrid parasites^7,8^ to the Sri Lankan isolates. The heterozygosity ratios were calculated as the portion of the heterozygous SNPs and indels across the entire genome in a sample (het/het+hom). Each dot represents a single SRA record. Isolates from Ethiopia previously identified as hybrid parasites^7,8^ and their respective frequency shown in blue. Sri Lankan isolates from the SL1, SL2 and SL3 groups are shown in red and their group assignment is highlighted.

To compare the overall level of heterozygosity in the Sri Lankan groups to other *L. donovani* strains, all 684 genomes used to generate the phylogeny tree in Figure 1 were aligned to the *L. donovani* reference genome sequence^24^ and variant sites were analysed using the VarScan2 software^32^. The frequency of heterozygous vs homozygous SNPs across the entire genome was calculated for each isolate to determine the level of heterozygosity for each *L. donovani* genome in the NCBI database. The ratios of heterozygous SNPs were plotted for all the sequenced *L. donovani* isolates and are shown in Figure 2B. A group of isolates originating from Ethiopia were considered separately as these were previously reported to be intra-species hybrids of two distinct *L. donovani* populations and therefore served as a benchmark for hybrid parasites^7,8,31^. The Ethiopian hybrid *L. donovani* isolates are highlighted in blue in Figure 2B. As shown in red, all isolates from the SL1 group fall within the normal distribution of heterozygous polymorphisms for *L. donovani*. All five isolates from the Sri Lanka SL2 group have a high ratio of heterozygous SNPs above the known hybrid group from Ethiopia. In comparison, all three isolates from the SL3 group are close to the overall distribution of *L. donovani* isolates. Due to the distance and the high heterozygous SNP frequency of the SL2 group compared to the entire *L. donovani* global population (Fig.1B, Fig.2B, Supp. Fig. S1), we investigated the possibility that these were interspecies hybrid parasites.

### Cutaneous disease-associated *Leishmania* species contribute hybrid parental genomes

Highly heterozygous SNPs were of interest because they could be derived from non-*L. donovani* species. To investigate this possibility, nucleotides corresponding to the site of each SNPs were altered to correspond to the non-*L. donovani* reference nucleotide to reconstruct the genes contributed by a potential non-*L. donovani* parent as outlined in Figure 3A. The reconstructed genomes were then compared to all Old World reference *Leishmania* strains available on TriTrypDB^33^ using BLAST. To validate this methodology, a previously reported hybrid parasite between *L. major* and *L. infantum* (IMT211) was used as a positive control test assay^11^ while the SL1 non-hybrid group previously characterized^24,25^ serves as an internal negative control. As shown in Figure 3B, the reconstructed genes using the non-reference nucleotides polymorphisms from the control sample (IMT211) matched almost entirely *L. major* confirming that this method of analysis can quantify the sequence contributions from the non-*L. donovani* parent at the whole genome level.

**Figure 3.**
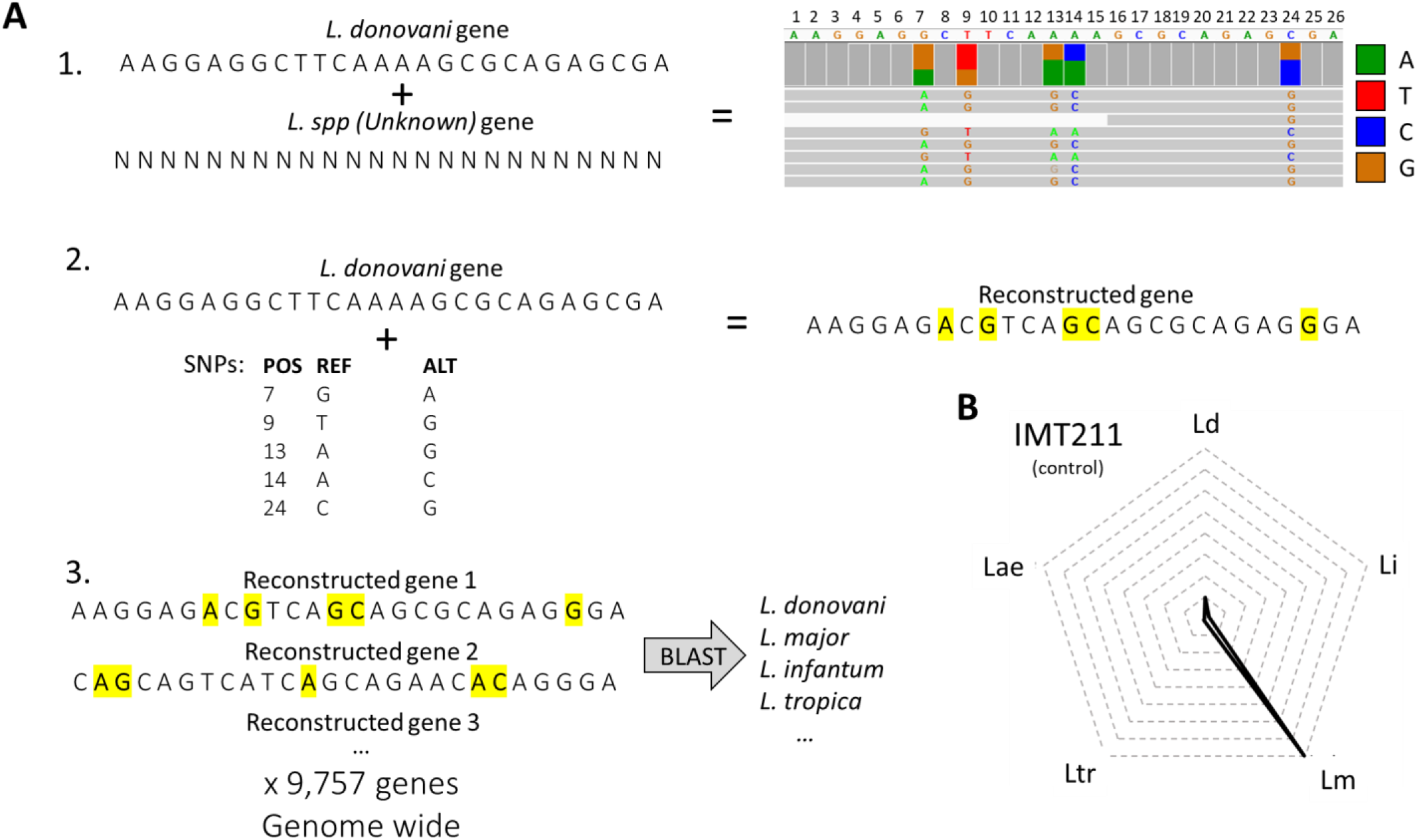
Methodology used to determine the parental strain lineage. **A.** Hybrid SNP loci were assumed to have received one allele from *L. donovani* and one allele from an unknown parent resulting in alignments with polymorphisms occurring at +/-50%. Gene sequences from the Sri Lanka reference *L. donovani*^24^ (REF) were transformed at the position (POS) of each SNP (ALT) across the entire genome to reconstruct the gene sequences of the unknown parent *Leishmania* species. All reconstructed gene sequences were then compared to a *Leishmania* database containing all Old World *Leishmania* reference genomes by BLAST searches and assigned an originating species and strain. **B.** Control analysis using a known hybrid (IMT211) with *L. major* and *L. infantum*^11^ showing reconstructed genes mostly matching *L. major*. Each level in the radar plot corresponds to 1,000 gene matches in the corresponding species (dotted lines).

Using the above method of analysis, the reconstructed genomes from the SL1 group matched almost exclusively to members of the *L. donovani* species complex^24^. Analysis of the reconstructed genomes from the highly heterogenous SL2 group however revealed that the SRR6257364 isolate had SNPs of *L. major* origin in almost half its genome and SRR6257365 contained SNPs of *L. major* origin in about 20% of its genome (Table 1, Figure 4A, SL2 A, red). Moreover, the reconstructed genes from samples SRR6257366, SRR6257367 and SRR6257369 matched mostly the *L. tropica* reference genome (Table 1, Figure 4A, SL2 B, green). In comparison, the SL3 group (Isolates SRR6257368, SRR6257370, SRR6257371) that clustered closer to the African strains (Fig. 1B), contained very few reconstructed gene matches outside of the *L. donovani* complex (*L. donovani/L. infantum*) (Table 1, Figure 4A, SL3, blue). For clarity, only the *Leishmania* species with gene matches are shown, the complete Old World species gene comparison is shown in Supplementary Fig S2.

**Table 1.** Old World *Leishmania* species genome matches after alternative allele gene reconstruction. Cumulative results of the best scoring species matches from genome wide BLAST searches using alternative allege gene reconstruction. In every sample, each gene across the genome was modified to reflect the sample polymorphism. These genes were then compared to the complete reference genomes of all Old World Leishmania species and the highest scoring alignment per gene was counted as one species match for that sample. Significant changes compared to the SL1 group are marked with *, **, *** or **** to denote p <0.05, p<0.01, p>0.001, p<0.0001 respectively based on 2-way ANOVA with multiple comparisons within each row.

|  | SL2 A |  | SL2 B |  |  | SL3 |  |  | SL1 |  |  |  |
| --- | --- | --- | --- | --- | --- | --- | --- | --- | --- | --- | --- | --- |
|  | SRR6257364 | SRR6257365 | SRR6257366 | SRR6257367 | SRR6257369 | SRR6257368 | SRR6257370 | SRR6257371 | ERR205734 | SRR7133731 | SRR7133732 | SRR7133733 |
| *L. donovani* | 3821**** | 4979**** | 662**** | 2871**** | 1711**** | 6492**** | 6495**** | 6475**** | 8012 | 8012 | 8011 | 8015 |
| *L. infantum* | 1770 | 2214 | 296 | 1404 | 860 | 2942**** | 2940**** | 2958**** | 1423 | 1423 | 1424 | 1420 |
| *L. major* | 3738**** | 2160**** | 230 | 203 | 227 | 47 | 46 | 50 | 46 | 46 | 46 | 46 |
| *L. tropica* | 93 | 115 | 7671**** | 4568**** | 5977**** | 112 | 112 | 110 | 111 | 111 | 111 | 111 |
| *L. aethiopica* | 6 | 6 | 412 | 317 | 496 | 0 | 0 | 0 | 0 | 0 | 0 | 0 |
| *L. arabica* | 1 | 2 | 8 | 5 | 5 | 0 | 0 | 0 | 0 | 0 | 0 | 0 |
| *L. gerbilli* | 33 | 21 | 66 | 33 | 69 | 0 | 0 | 0 | 0 | 0 | 0 | 0 |
| *L. enriettii* | 0 | 0 | 0 | 0 | 0 | 0 | 0 | 0 | 0 | 0 | 0 | 0 |
| *L. tarentolae* | 7 | 6 | 16 | 15 | 16 | 6 | 6 | 6 | 6 | 6 | 6 | 6 |
| *L. turanica* | 288 | 254 | 396 | 341 | 396 | 158 | 158 | 158 | 159 | 159 | 159 | 159 |

**Figure 4.**
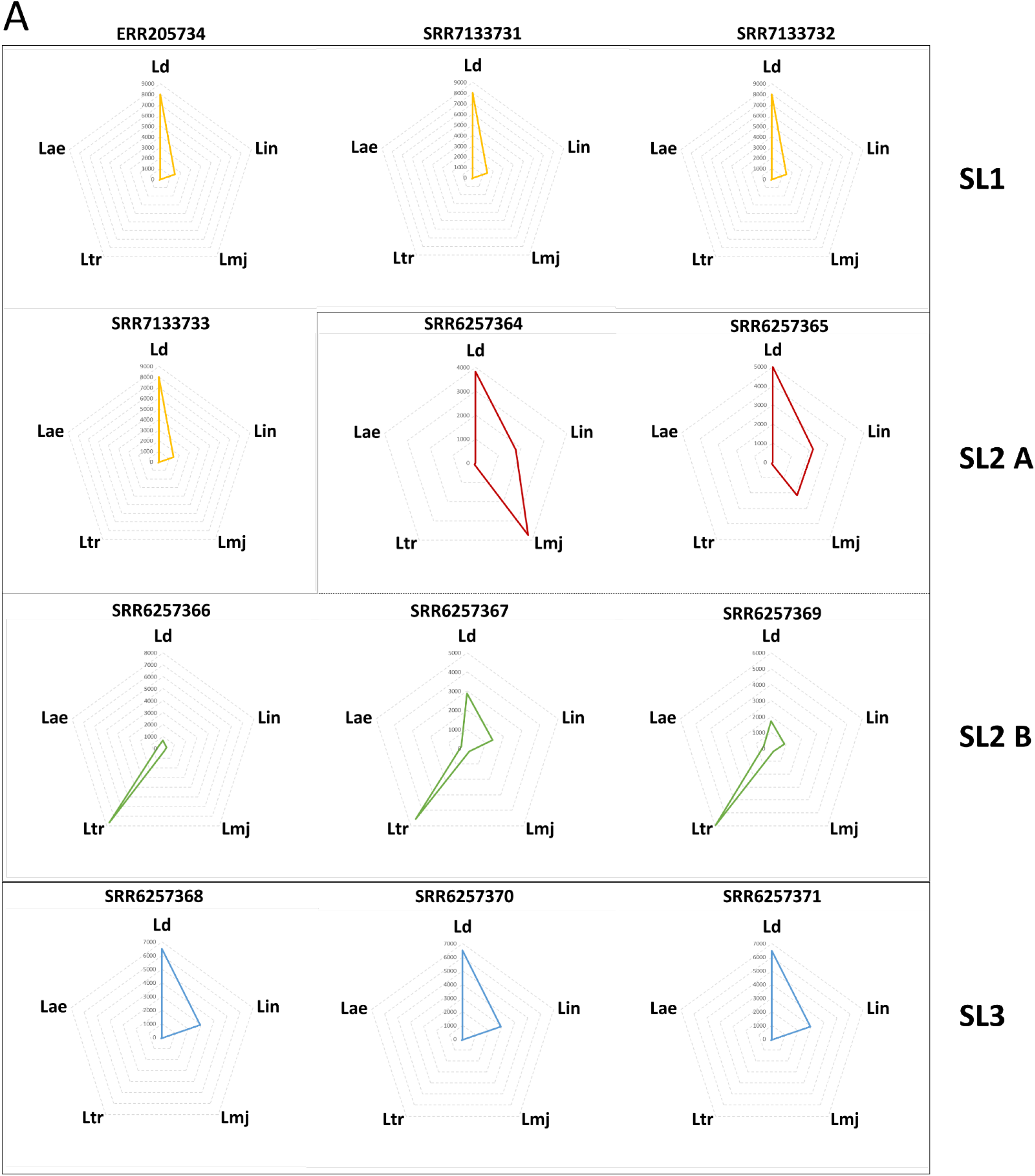
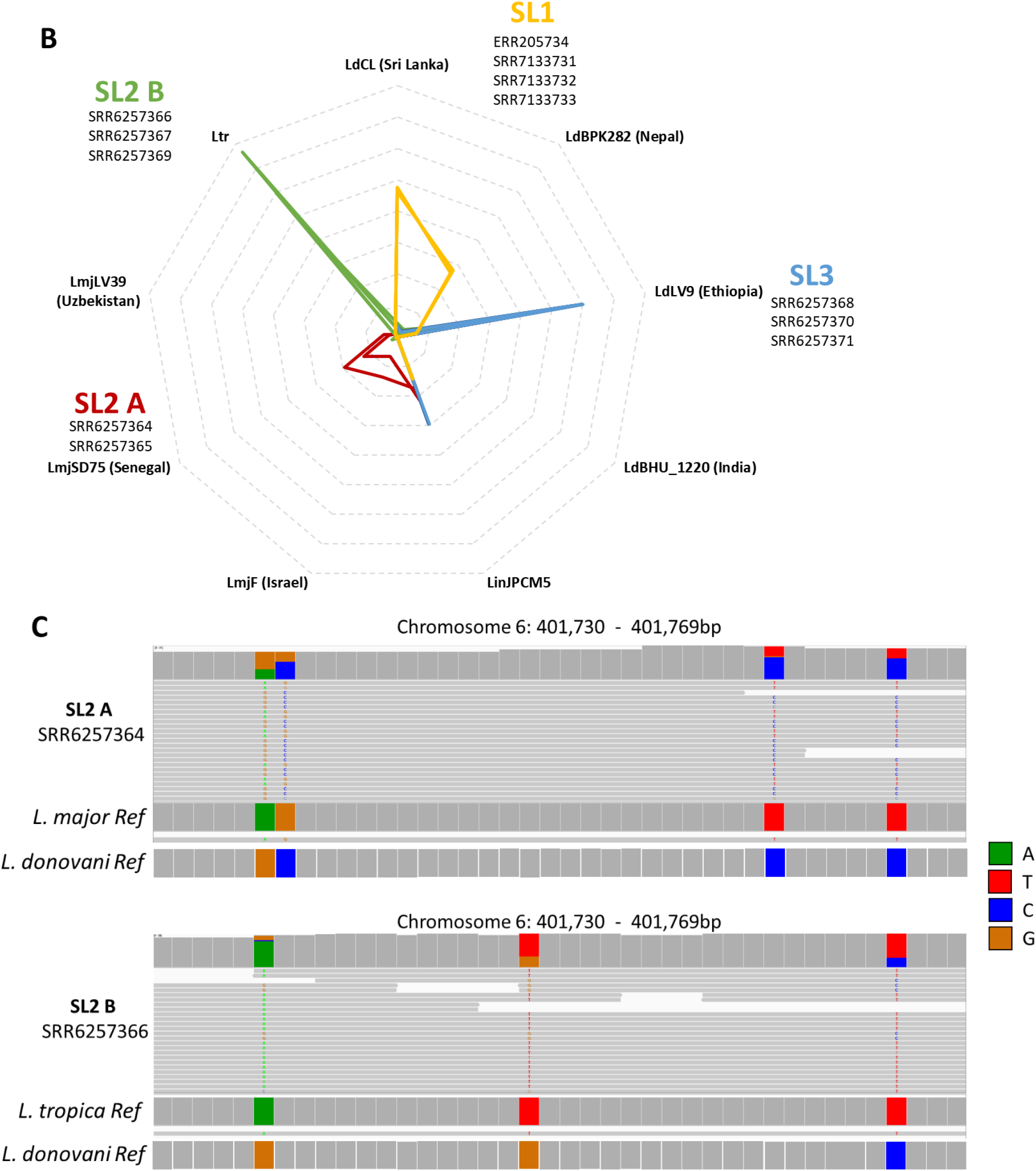
Evidence of hybrid genotypes with *L. major* and *L. tropica*. **A.** Distribution of species origin of reconstructed genes as determined by BLAST analysis. Reconstructed genes from all SL1 isolates match nearly exclusively the *L. donovani* species complex. Two SL2 isolates match predominantly to *L. donovani* complex species but retained genes from *L. major* (SL2 A, top). Three SL2 isolates match with *L. tropica* more than with the *L. donovani* complex (SL2 B, middle). Three isolates match the *L. donovani* complex with almost no match outside *L. donovani* (SL3, bottom). **B.** Distribution of the origin of reconstructed genes to different reference strain genomes as determined by BLAST analysis. Each level in the radar plots corresponds to 1,000 gene matches in the corresponding species (dotted lines). All SL1 isolates show a greater match to Sri Lankan and Nepalese reference genomes (Orange). The two SL2 A subgroup isolates show a greater match with the *L. major* strain SD75 from Senegal than with the Friedlin strain from Israel or LV39 strain from Uzbekistan (red). The three SL2 B subgroup isolates show a single match with *L. tropica* (green). The three SL3 group isolates show a preferential match to the African LV9 strain of *L. donovani* (blue). **C.** Representative alignment of a *L. major* and *L. tropica* hybrid on the same section of chromosome 6 showing heterozygous polymorphism matching *L. major* and *L. tropica* respectively.

The above analysis identified genes originating from different reference *Leishmania* species. We next used this methodology to attempt to further narrow the origins of the hybrid genes present in the Sri Lankan isolates. As shown in Figure 4B and Table 2, the SL1 group matched mostly with the reference sequences from Sri Lankan and Nepal, while the non-*L. donovani* parent from the *L. major* hybrids classified as group SL2 A (SRR6257364, SRR6257365) was more related to *L. major* strain SD75 (isolated from Senegal) than to *L. major* strain LV39 (Isolated from Uzbekistan) or the reference *L. major* Friedlin strain (isolated from Israel). As there is only one reference *L. tropica* strain, samples SRR6257366, SRR6257367 and SRR6257369 (SL2 B) all clustered to this one reference. With respect to the SL3 group, the reconstructed genes from SRR6257368, SRR6257370 and SRR6257371 were almost exclusively derived from the African LdLV9 reference strain^34^.

**Table 2.**
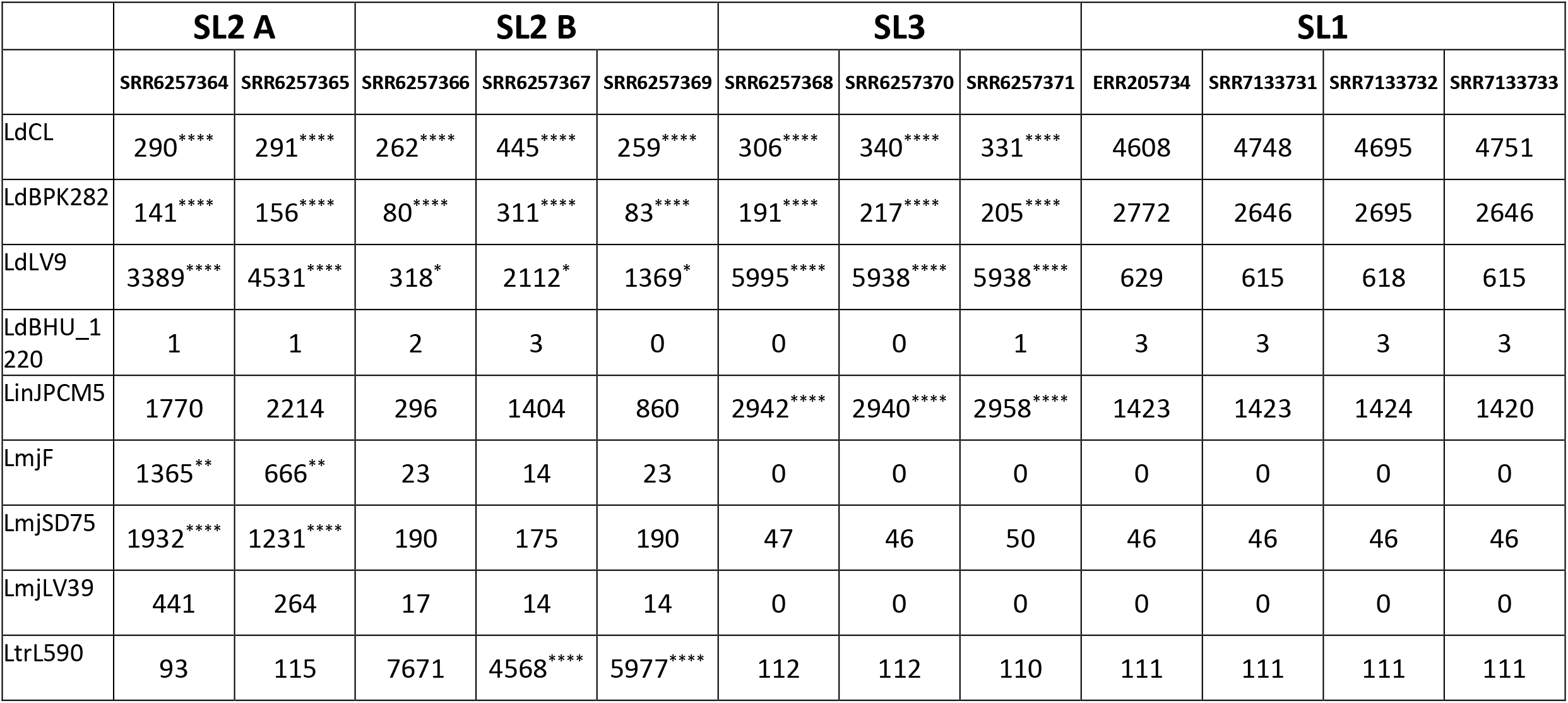
Old World *Leishmania* strains genome matches after alternative allele gene reconstruction. Cumulative results of the best scoring strain matches from genome wide BLAST searches using alternative allege gene reconstruction. In every sample, each gene across the genome was modified to reflect the sample polymorphism. These genes were then compared to the complete reference genomes of all Old World Leishmania strains and the highest scoring alignment per gene was counted as one strain match for that sample. Significant changes compared to the SL1 group are marked with *, **, *** or **** to denote p <0.05, p<0.01, p>0.001, p<0.0001 respectively based on 2-way ANOVA with multiple comparisons within each row.

|  | SL2 A |  | SL2 B |  |  | SL3 |  |  | SL1 |  |  |  |
| --- | --- | --- | --- | --- | --- | --- | --- | --- | --- | --- | --- | --- |
|  | SRR6257364 | SRR6257365 | SRR6257366 | SRR6257367 | SRR6257369 | SRR6257368 | SRR6257370 | SRR6257371 | ERR205734 | SRR7133731 | SRR7133732 | SRR7133733 |
| LdCL | 290**** | 291**** | 262**** | 445**** | 259**** | 306**** | 340**** | 331**** | 4608 | 4748 | 4695 | 4751 |
| LdBPK282 | 141**** | 156**** | 80**** | 311**** | 83**** | 191**** | 217**** | 205**** | 2772 | 2646 | 2695 | 2646 |
| LdLV9 | 3389**** | 4531**** | 318* | 2112* | 1369* | 5995**** | 5938**** | 5938**** | 629 | 615 | 618 | 615 |
| LdBHU_1<br>220 | 1 | 1 | 2 | 3 | 0 | 0 | 0 | 1 | 3 | 3 | 3 | 3 |
| LinJPCM5 | 1770 | 2214 | 296 | 1404 | 860 | 2942**** | 2940**** | 2958**** | 1423 | 1423 | 1424 | 1420 |
| LmjF | 1365** | 666** | 23 | 14 | 23 | 0 | 0 | 0 | 0 | 0 | 0 | 0 |
| LmjSD75 | 1932**** | 1231**** | 190 | 175 | 190 | 47 | 46 | 50 | 46 | 46 | 46 | 46 |
| LmjLV39 | 441 | 264 | 17 | 14 | 14 | 0 | 0 | 0 | 0 | 0 | 0 | 0 |
| LtrL590 | 93 | 115 | 7671 | 4568**** | 5977**** | 112 | 112 | 110 | 111 | 111 | 111 | 111 |

As *L. tropica* and *L. major* are genetically closely related, we manually inspected some of the generated alignments to confirm that it was possible to accurately assign the reconstructed genes as belonging to either *L. tropica* or *L. major*. As in the representative alignment in Figure 4C, the polymorphisms between the *L. tropica* and *L. major* hybrids were frequent enough to be discriminatory. As *L. major* and *L. tropica* parasites are not present in Sri Lanka, and considering the phylogenetic tree shown in Figure 1B, it is likely that the hybrids originated in East Africa and subsequently imported into Sri Lanka from infected individuals.

To verify the possibility that SL2 groups contained genetic material originating from *L. major* and *L. tropica*, we performed an additional genetic comparison and phylogeny analysis including sequences from *L. major* and *L. tropica*. As shown in Supplementary Figure S3, all SL2 isolates are placed at various distances along the same branch as *L. major* and *L. tropica*. Further, this branch containing the putative inter-species hybrids and cutaneous species is shown to originate from the African *L. donovani* lineages.

We also investigated whether it was possible to separate the haplotypes by separating the reads originating from chromosomes with different species origins through read-based phasing to further verify the identification of species by the BLAST analysis described above. This was however complicated by chimeric phase sets, likely due to insufficient coverage. As shown in Supplementary Figure S4A, SNP dense regions originating from the *L. major* parent of the hybrid phased in this alignment are assigned alternatively to the different phases rather than remaining continuous. Nevertheless, phasing of a short segment on chromosome 1 followed by phylogenetic analysis was consistent with the BLAST comparison approach as outlined in Figure 3A. As shown in Supplementary Figure S4B, the haplotypes originating from the SL2A and SL2B isolates clustered on opposite branches between *L. donovani/L. major* and *L. donovani/L. tropica* respectively, consistent with the BLAST analysis of the origin of the non-*L. donovani* hybrid parent.

### Chromosomal recombination in *L. donovani* hybrid strains

We next investigated whether there was evidence for recombination between *L. donovani* and *L. major* homologous chromosomes. As shown in a representative alignment of the two SL2A *L. donovani/L. major* hybrids for the same section of chromosome 19, blocks of nucleotides consisting of homozygous *L. donovani* sequences and heterozygous *L. donovani/L. major* sequences were evident (Figure 5A). This pattern of mixed or single parental origin sequences can be seen for the SL2 isolates throughout all the chromosomes as shown in the example of chromosome 36 (Figure 5B) and the whole genome (Supplementary Fig S5).

**Figure 5.**
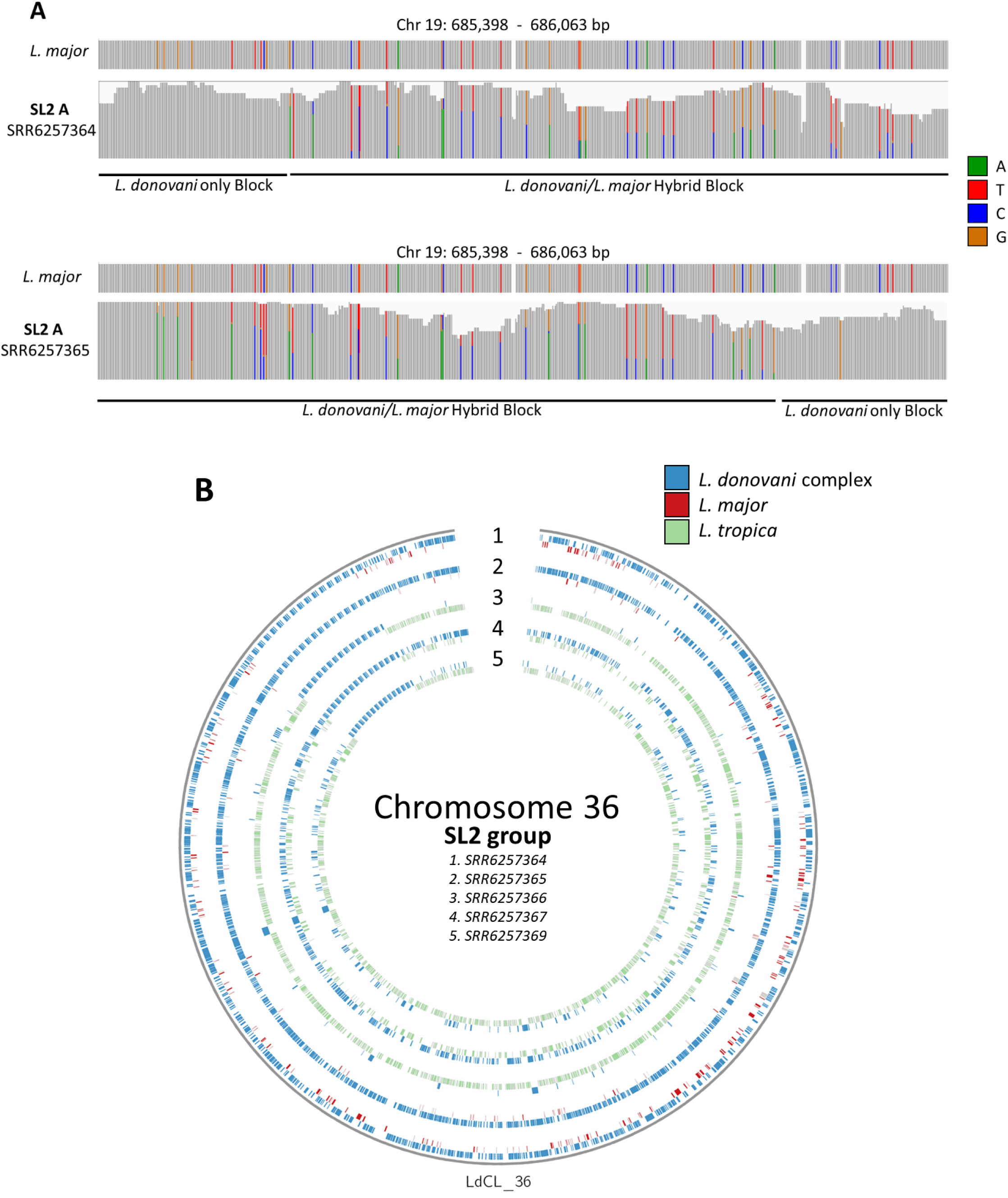
Chromosomal recombination in *L. donovani* hybrid strains with *L. major* and *L. tropica*. **A.** Representative alignment of genomes from two *L. major* hybrid parasites on the same section of chromosome 19 showing short length blocks of single (*L. donovani* only) or mixed parent ancestry (*L. donovani/L. major* hybrid). **B.** Representation of chromosome 36 in all isolates in the SL2 group. Each marker represents a single gene. Genes of *L. donovani* species complex origin are marked in blue. Genes with hybrid ancestry (*L. major & L. donovani*, or *L. tropica & L. donovani*) are colored in red and green respectively.

### *L. donovani* isolates with low heterozygosity (SL3 group) contain *L. major* SNPs and has conserved aneuploidy

While the five SL2 isolates were highly heterozygous with a large fraction of SNPs matching the *L. tropica* or *L. major* genomes, the three SL3 isolates contained a low level of heterozygosity (Fig. 2B). We therefore investigated whether other genomic alterations were present that could explain the common phenotype of these variants. Upon close inspection, these isolates also contained some short regions with polymorphisms in common with *L. major*. In contrast, the sequences from the SL1 isolates contained no detectable polymorphisms in common with *L. major*. Although most of the polymorphisms in the SL3 group were no longer present at 50% allele frequency (diploid heterozygous), many of the polymorphisms were in common with the SL2 group as shown in this representative section of chromosome 6 (Figure 6A). Further, some of the SNPs are retained without a loss of allele frequency to match the *L. major* sequence. Supplementary Table S2 contains a list of genes with *L. major* non-synonymous polymorphisms retained in all samples of the SL3 group.

**Figure 6.**
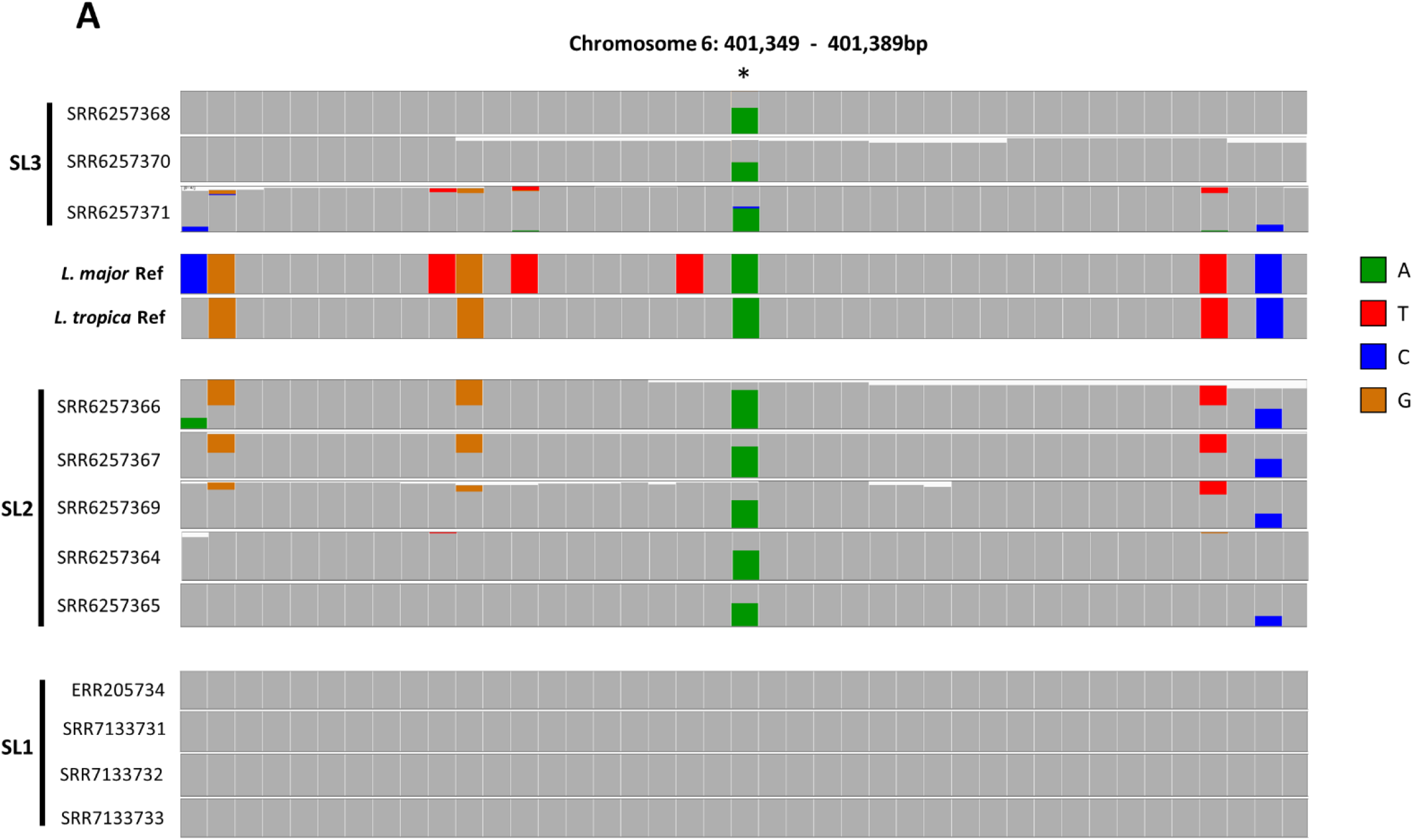
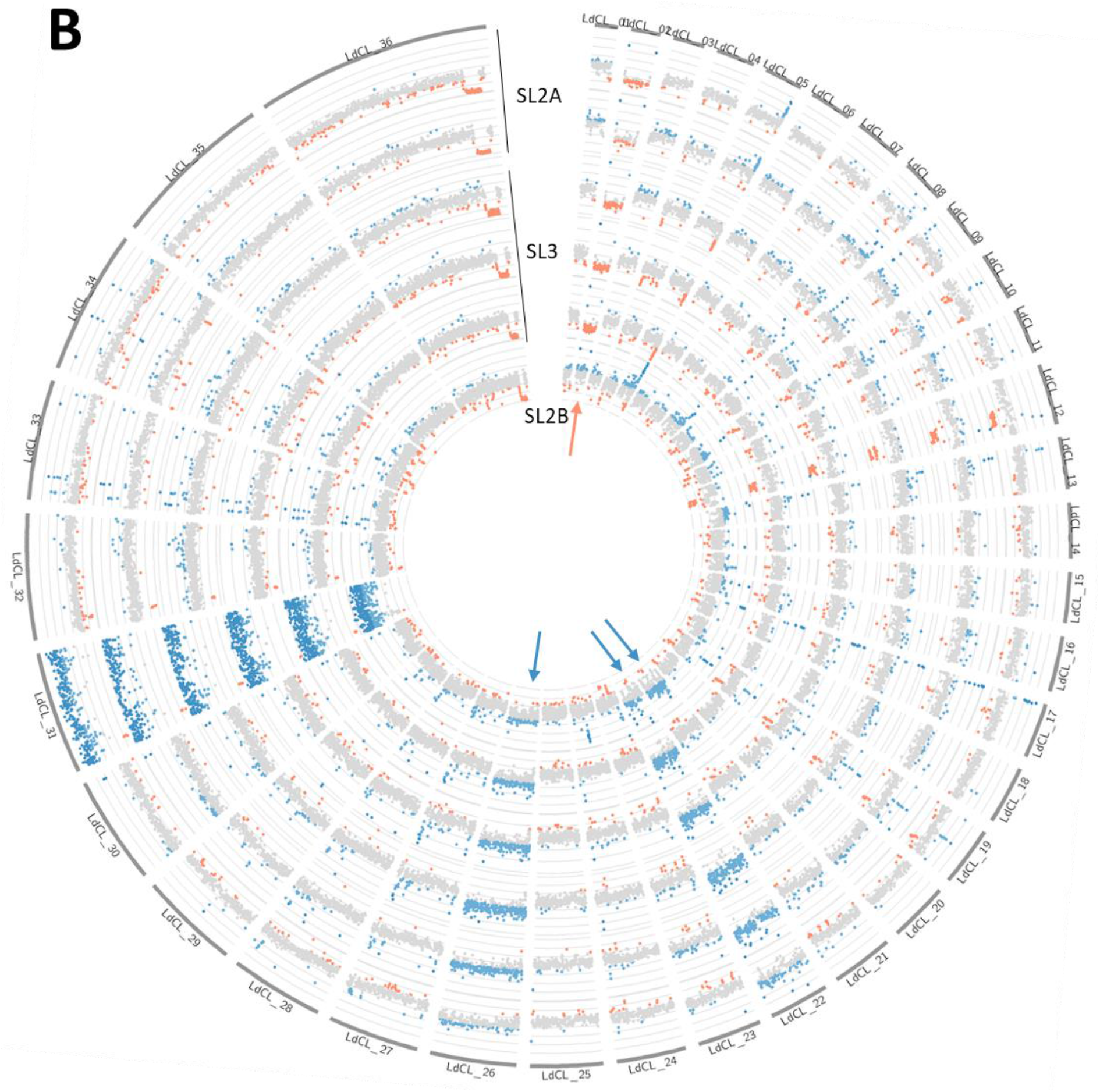
Evidence of ancient hybridization in SL3 group samples. **A.** Prevalence of polymorphisms across all Sri Lanka isolates on a portion of chromosome 6 compared with the reference *L. major* and *L. tropica* sequences. Grey boxes represent Sri Lanka reference *L. donovani* nucleotides. The SL3 group have retained varied levels of *L. major* alleles depending on the sample (upper 3 alignments). The SL2 group retained *L. major* or *L. tropica* polymorphisms. The SL1 group does not share polymorphisms with either *L. tropica* or *L. major*. The polymorphism highlighted in the center (*) of the alignment and retained in all SL2 and SL3 isolates results in a Cys267Tyr change in the LdCL_060014600 gene and matches the *L. major* allele, shows that some polymorphisms appear to be more stable in the hybrid genomes. **B.** Conservation of aneuploidy patterns across SL3 and SL2A group samples. Chromosomal coverage was determined and colored according to mean coverage (grey), decrease coverage (red) or increase coverage (blue) in sequencing depth. Coverage at each gene location across the entire genome shows the *L. major* hybrid isolates (SL2A) have reduced average copies of chromosome 2, and increased copy number of chromosome 22 and 26. The SL3 isolates show the same aneuploidy pattern as the *L. major* hybrids (SL2A). In comparison, SL2B *L. tropica* hybrid parasites have normal coverage at chromosome 2, a unique increase at chromosome 21, and share the increase across chromosomes 22 and 26. All isolates appear diploid for all other chromosomes with the exception of naturally tetraploid chromosome 31.

Modulation of aneuploidy and unequal crossing over has been shown to be an evolutionary mechanism employed by *Leishmania*^35,36^. As the SL3 group showed evidence of ancient hybridization with *L. major* sharing polymorphic sites with the SL2 group (Figure 6A), we therefore investigated whether the SL2 and SL3 groups shared other features including conserved aneuploidy across the hybrid subgroups. Genome sequencing coverage was compared to determine the chromosome copy numbers. As shown in Figure 6B, the SL2A (*L. major/L. donovani* hybrids) and SL3 (*L. major/L. donovani* ancient hybrids) isolates have a conserved aneuploidy pattern consisting of a decreased copy number of chromosome 2 (red arrow) and an increased copy number of chromosomes 22 and 26 (blue arrows). Note also that chromosome 31 is tetraploid in all *Leishmania* species and serves here as a reference for an increased copy number chromosome. The SL2B isolate (*L. tropica/L. donovani* hybrids) shown in Figure 6B (and also in Supplementary Figure S5) does not share the decreased copy number for chromosome 2 and appears to have a unique increase in copy number of chromosome 21 and an increase of chromosomes 22 and 26 shared with the SL2A and SL3 groups. For comparison to SL2A and SL3, only one SL2B isolate is shown in Figure 6B and all three SL2B isolates are consistent and shown in Supplementary Figure S5. Overall, these observations show that there are similarities between the SL2A and SL3 groups with respect to conservation of polymorphisms and pattern of aneuploidy.

## Discussion

This study presents evidence that hybrid *L. donovani/L. major* and *L. donovani/L. tropica* parasites are associated with cutaneous leishmaniasis in Sri Lanka. Although this is an important observation, it is also remarkable that multiple atypical *L. donovani* hybrids (SL2 and SL3 groups) and non-hybrids (SL1)^24,25^ associated with cutaneous leishmaniasis culminate in Sri Lanka concomitant with the exclusion of visceral leishmaniasis that is now virtually non-existent^20^. As *L. major* and *L. tropica* parasites are not present in Sri Lanka, and considering the phylogenetic trees shown in Figure 1B and Supplementary Fig S3, it is likely that the hybrids originated in East Africa and were subsequently imported into Sri Lanka from infected individuals. Once in Sri Lanka, there appears to be an environmental selection for propagation of these atypical *L. donovani* parasites that cause cutaneous leishmaniasis in the local population.

It will be interesting to investigate the vector and potential non-human reservoirs that selects for such atypical *L. donovani* strains since virtually all human cutaneous leishmaniasis causing species outside of Sri Lanka have an animal reservoir. Notably, the probable vector *L. donovani* in Sri Lanka is *P. argentipes* subspecies *glaucus* that has a preference for animal rather than human blood. This differs from *P. argentipes* subspecies *sensu lato* the vector for *L. donovani* in India, that is anthropophilic^37^. Further, this is also different from the *P. orientalis* and *P. alexandri* vectors of *L. donovani* in Africa^7^ and as inter-species hybridization has previously been shown to confer increased vector competence to hybrid parasites ^38^, the hybrids described herein could benefit from a wider permissive vector repertoire.

Starting from an original outcross hybridization with *L. donovani*, subsequent replication through mitosis or a combination of mitosis and meiosis involving cross over events potentially resulted in the retention of genetic information from *L. major* or *L. tropica*. This is consistent with the observation that recombination can occur in the progeny of *L. donovani* complex parasites as was demonstrated for the *L. infantum* and *L. donovani* hybrid associated with cutaneous leishmaniasis in Turkey^5^.

These observations are also in agreement with the proposed model of mosaic aneuploidy^35^ recently supported by whole genome sequencing data^36^ where *Leishmania* can discard deleterious or retain beneficial alleles. Indeed, recent research shows the genome of *Leishmania* is highly dynamic during replication resulting in a high genomic diversity across the pooled population^35,36^. Through this process, progeny with a combination of *L. major* or *L. tropica* alleles with *L. donovani* alleles can arise if they posses a fitness advantage for propagation in a particular environment.

Whole genome sequence analysis of *L. donovani* causing cutaneous leishmaniasis strains originally isolated from Sri Lanka were however not hybrid parasites and contain 83 gene mutations including altered A2 virulence genes and a polymorphism in the RagC gene from the mTOR signalling pathway^24,25,39^. None of these originally identified 83 genes mutations were present in the more recently sequenced *L. donovani* strains from Sri Lanka included in this analysis (GenBank ID PRJNA413320)^26^. This argues that there are diverse mechanisms for *L. donovani* to lose virulence for causing visceral disease in Sri Lanka and become associated with the less virulent cutaneous disease and these include genetic changes within the *L. donovani* genome^24, 25^ and potentially as demonstrated within, propagation of *L. donovani* hybrid strains with *L. major* and *L. tropica*.

Parasites were also identified that contained relatively small amounts of hybrid gene polymorphisms in the SL3 group suggesting that introgression was more ancient in SL3 than SL2 resulting in fewer polymorphic alleles. This is essentially a natural selection experiment where heterozygosity is reduced by the removal of SNPs that are not beneficial and the retention of SNPs that result in more fit parasites for the Sri Lankan environment. Supplementary Table S2 contains those genes that have retained *L. major* non-synonymous SNPs despite having low levels of heterozygosity. Some of these genes could have been selectively retained during propagation in Sri Lanka and functional analysis of these genes could identify their role in disease tropism. Nevertheless, none of the genes in Supplementary Table S2 are in common with the mutant genes from the non-hybrid SL1 group^24,25^ consistent with the argument that there are multiple mechanisms for genotypic changes in *L. donovani* to mediate cutaneous leishmaniasis.

Interestingly, while the patterns of polymorphisms appear varied across the isolates, the aneuploidy seen in the SL2 and SL3 parasites appears to be more conserved (Fig. 6A, B) suggesting an external pressure is driving those parasites away from a normal diploid genome. Polymorphisms identified above may be further narrowed by prioritizing polymorphisms retained on chromosomes presenting an unusual ploidy pattern.

The evidence presented herein for the existence of 4 different populations of *L. donovani* parasites in Sri Lanka (SL1, SL2A, SL2B and SL3) could help reconciliate differences in lesion morphology^40,41^, spatial distribution^20^ and drug susceptibility^26^ previously reported across Sri Lanka. Indeed, parasites causing lesions that share features of *L major* or *L. tropica* infection^40^ or with variable tolerance to sodium stibogluconate^26^ could consist of different *L. donovani* hybrids with varying amount of genes from either *L. major* or *L. tropica* as they have different pathological features^42^ and drug sensitivity^43^. Further, these co-existing populations supports both theories of recent introduction or prolonged existence of endemic *L. donovani* parasites in Sri Lanka^20^. It would be interesting to compare the pathology caused by SL2 and SL3 group parasites or the *L. major* vs *L. tropica* hybrids as these different parasites could be responsible for the different cutaneous disease profiles identified between the North and South parts of the island^40^.

Genetic comparison of the genome of *L. donovani* origin in those hybrids is most similar to the LV9 strain of Ethiopian origin^34^ rather than the reference Sri Lankan or Indian/Nepalese sequences while also assigning the genomic sequences of *L. major* origin to a Senegalese reference (Table 2, Fig. 1B, Supplementary Fig S3, Fig. 4B). *L. donovani* parasites from Ethiopia have previously formed intra-species hybrid parasites between the divergent northern and southern populations^6–9^. Further, studies on *L. tropica* have identified intra-species hybridization^44^ and *L. major* hybrids have previously been isolated^11^ indicating these two parasite species are amenable to the generation of hybrid progeny. This study however provides novel evidence through whole genome analysis for the generation of natural *L. donovani* hybrids with *L. major* and *L. tropica*. Both *L. tropica* and *L. major* are present in Ethiopia in similar animal populations along with their sandfly vectors^45^. Not only is *L. tropica* present in Ethiopia, but it was also reported to share similar possible animal reservoirs with *L. donovani* in Ethiopia^46^. Taken together, it is feasible that *L. donovani* parasites from Ethiopia could generate hybrid parasites with *L. tropica* and *L. major*. These hybrids could have entered Sri Lanka as suggested by the grouping of SL3 parasites in the North-Ethiopian/Sudanese cluster (Fig. 1B, Supplementary Fig S3).

Sri Lankan military personnel are routinely deployed in *Leishmania*-endemic countries as part of UN peace keeping missions and returning soldiers are a known risk factor for importing parasitic diseases^47^. Indeed, chemoprophylaxis and screening against Malaria upon returning from a mission is routine as part of Sri Lanka personnel deployments in countries such as Sudan^47^. As argued within this and other studies^24–26^, the genetic evidence argues there have been multiple sources of *L. donovani* entry into Sri Lanka, yet there is no visceral disease and atypical cutaneous leishmaniasis has successfully propagated^20^. Future studies must address why atypical *L. donovani* parasites would be imported and propagated in Sri Lanka instead of visceral disease-causing *L. donovani* which are vastly more common in neighboring India and Africa.

## Methods

### Data collection

All sequencing information used in our analysis was obtained from publicly available records on the Sequencing Reads Archive (SRA) repository provided by NCBI^27^. A search was performed to obtain all sequencing records for the *Leishmania donovani* organism (NCBI Taxonomy ID 5661). The read sets were filtered to remove any data originating from unknown or transcriptomic cDNA source. The remainder of gDNA reads sets were filtered to remove targeted capture experiments and retained only whole genome random selection libraries. As long reads are error prone and more suitable for assembly than SNP calling, non-Illumina long read sets were removed. To increase the accuracy of the SNP calls, only reads in paired mode were retained. Any SRA belonging to the BioProject PRJEB8793 were removed as they originate from single cell data and therefore did not contain enough coverage. Other samples with low coverage were removed manually after inspection of the alignments, the retained read sets used for phylogenetic analysis are listed in Supplementary Table S1. Samples with the ‘LK’ WHO country code (Sri Lanka) in their isolate name were grouped with our previous Sri Lankan data.

Sequences from NCBI BioProject PRJNA413320 were from 6 cutaneous leishmaniasis case isolates (SRR6257369, SRR6257364, SRR6257365, SRR6257368, SRR6257370, SRR6257371) and 2 isolates from visceral leishmaniasis patients with Leukemia and Diabetes Mellitus co-morbidities (SRR6257366 & SRR6257367) ^26^. As there have been fewer than 7 visceral leishmaniasis cases since 2004 in Sri Lanka and the majority of these having co-morbidities^21^, we have made the assumption that the SRR6257366 and SRR6257367 isolates do not cause active transmission of visceral leishmaniasis but likely have contributed to the 15000 reported cases of cutaneous leishmaniasis in Sri Lanka since 2001^19,20^.

### Alignment of all sequenced *L. donovani* isolates to the reference genome

The filtered 684 sequencing samples were distributed evenly across three Compute Canada clusters, Beluga at Calcul Quebec, Niagara at SciNet, and Cedar at WestGrid. The raw reads for each sample were downloaded using the SRA-Toolkit provided by NCBI^27^ using the ‘fastq-dump–split-files’ command. The alignment of all reads obtained from the SRA was performed as previously described^24^. Briefly, Illumina paired reads were aligned to the reference Sri Lanka genome^24^ using the Burrows-Wheeler Aligner^48^, file formats transformed using samtools^49^, and variant calling was done with VarScan2^32^ to generate VCF files with an alternate allele frequency cut-off of 20%. This pipeline was automated using in house scripts to process the data in parallel across 684 nodes on the clusters resulting in one VCF file per sample containing a list of all polymorphisms and their respective frequencies.

### *L. donovani* global strains phylogeny

The resulting 684 VCF files generated at the alignment stage described above were merged using BCFtools^49^ with the command ‘bcftools merge –missing-to-ref’ resulting in a single VCF file containg all the possible genetic polymorphisms identified in all 684 samples. The merged VCF file was imported into TASSEL version 5.0^50^ and subjected to a relatedness analysis to generate a distance matrix and a phylogenetic tree using the Neighbor-Joining algorithm. The phylogenetic tree was exported in Newick format and visualized using the Interactive Tree of Life (IToL)^51^ to assign colors to nodes and clades.

After identification of likely inter-species hybridization, additional samples from *L. major* and *L. tropica* whole genome sequencing experiments were aligned to the *L. donovani* LdCL genome to generate a list of polymorphisms between the species and these were added to the global *L*. *donovani* polymorphism VCF file and processed as above to generate a phylogenetic tree to place the putative hybrid strains.

### Heterozygosity of *L. donovani* isolates worldwide

Variant loci from the VarScan2 annotated VCF files were assigned a HET or HOM value based on the variant allele read frequency for each site for each sample if the software determined the SNP to be heterozygous or homozygous respectively. The frequency of HET and HOM variant annotations across the entire genome was calculated on a per sample basis resulting in a single data point per sample with the Heterozygous ratio defined as (Het/Het+Hom). As the SL1 isolates fall within the reference cluster, resulting in homozygous polymorphisms being masked by the reference which artificially shifts the Heterozygous ratio, these isolates were aligned to the Nepalese reference genome for the purpose of this calculation. The Sri Lankan isolates from BioProject PRJNA413320 (belonging to the SL2 and SL3 groups), our previous studies^24,25^, and one sample from PRJEB2600 (belonging to the SL1 group)were grouped together. 10 samples previously characterized as intra-species hybrids of *L. donovani* between the North and South clusters of Ethiopian *L. donovani*^7^ were grouped together as a hybrid sample positive control group. The remaining 662 samples were grouped together to represent the natural *L. donovani* distribution.

### Identification of species in the hybrid parasites

The genomic annotation from the LdCL strain previously generated^24^ were used to generate a region list of the genomic coordinates of every gene in the ‘chr:start-stop’ format. This region list was used as an input to samtools with the ‘faidx’ command to extract the genomic sequence corresponding to each gene locus from the LdCL reference genome file.

For each sample independently, the VCF file containing the location and nucleotide change of every polymorphism along the genome for that sample was used as an input file for BCFtools^52^ with the command ‘consensus’ to transform the genomic sequences from the reference generated in the previous step into their alternate or reconstructed sequences to reflect the genotype of each sample for 9,757 gene loci.

The complete genomes of all *Leishmania* species reference strains were downloaded from TriTrypDB v46^33^ (*L. major* Friedlin, *L. donovani* BPK282, *L. tropica* L590, *L. tarentolae* Parrot-TarII, *L. turanica* LEM423, *L.gerbilli* LEM452, *L. enriettii* LEM3045, *L. arabica* LEM1108 & *L. aethiopica* L147) in FASTA format. The genomes were concatenated into a single FASTA file and used as an input for NCBI BLAST+ v2.7.1 using the ‘command makeblastdb’ and ‘-dbtype nucl’ option to create a database of Old World *Leishmania* genomic sequences. For the refined search, all additional non-reference strains genetic sequences were later added (*L. donovani* LV9, *L. donovani* LdCL, *L. donovani* BHU1220, *L. major* SD75, *L. major* LV39).

The alternate or reconstructed sequences for each sample were then used as a list of input queries for this database using the command ‘blastn’ with the options ‘-max_target_seqs 1 -max_hsps 1 - outfmt “6 qseqid qcovs pident stitle” to create a report of 9,757 species or strain matches for each queried sample. These reports were then tabulated to generate radar plots depicting the probable genetic contributions of each species/strain per sample and the genomic coordinates used to generate chromosomal maps of parental blocks using circos^53^ to paint the regions on a circular karyotype representation of the *L. donovani* genome.

### Haplotype phasing

Read-based phasing was applied to separate the two putative haplotypes present in samples with high heterozygosity. For each sample, the BAM alignment file generated by the burrows wheeler aligner^48^ was additionally processed prior to polymorphism analysis by VarScan v2^32^ as described above. The alignment was processed using the ‘phase’ command from samtools^49^ with the option ‘-A’ to drop reads with an ambiguous phase resulting in two separate alignment files with segregated reads based on haplotype blocks. The two alignments files containing roughly half of the original reads each were manually inspected for concordance along the coordinates output by the tools as being continuous phase blocks by manual inspection. Alternate gene reconstruction was performed on each phase of each sample independently using BCFtools^52^ as described above and aligned with orthologous reference *Leishmania* sequences using Clustal Omega^54^ to generate a phylogenetic tree of the phased samples.

## Supporting information

Blast Results

## Role of the funding source

This research was support by grant from the Canadian Institutes of Health Research (CIHR) to GM and a doctoral training award from the Fond de Recherche du Quebec en Santé (FRQS) to PL. The funding agencies had no role in the design, collection, analysis, or decision to publish this study.

## Declaration of interests

The authors declare no financial or personal competing interest.

## Contributions

PL designed the study, collected and analysed the data and wrote the manuscript. GM helped design the study, wrote and edited the manuscript. Both authors have read and approved the final version of the manuscript.

## Acknowledgements

We thank Wen Wei Zhang, Kayla Paulini, and Jesse Shapiro for their insightful feedback and suggestions. All DNA sample collection and sequencing were performed by their respective research groups, for complete attribution refer to the corresponding SRA accession page.

This research was enabled in part by support provided by Calcul Quebec (Beluga cluster, https://www.calculquebec.ca/en/), SciNet (Niagara cluster, https://www.scinethpc.ca/niagara/), WestGrid (Cedar Cluster, https://www.westgrid.ca/) and Compute Canada (www.computecanada.ca).

## Data availability

### Accession Numbers

All data used in this study were obtained from publicly available sources, all accession numbers used are listed in Supplementary Table S1.

### Data files

The phylogenetic tree generated in this study is available in an interactive format provided by iToL at: https://itol.embl.de/tree/1322162673368791580134755

### Code availability

All software and methodologies used are described within Methods.

## Notes

### Competing Interest Statement

The authors have declared no competing interest.

## References

1 WHO. Leishmaniasis. https://www.who.int/leishmaniasis/burden/en/.

2 Burza S, Croft SL, Boelaert M. Leishmaniasis. Lancet 2018; 392: 951–70.

3 Rijal S, Sundar S, Mondal D, Das P, Alvar J, Boelaert M. Eliminating visceral leishmaniasis in South Asia: The road ahead. BMJ. 2019. DOI:10.1136/bmj.k5224.

4 Scorza BM, Carvalho EM, Wilson ME. Cutaneous manifestations of human and murine leishmaniasis. Int. J. Mol. Sci. 2017. DOI:10.3390/ijms18061296.

5 Rogers MB, Downing T, Smith BA, et al. Genomic Confirmation of Hybridisation and Recent Inbreeding in a Vector-Isolated Leishmania Population. PLoS Genet 2014. DOI:10.1371/journal.pgen.1004092.

6 Zackay A, Nasereddin A, Takele Y, et al. Polymorphism in the HASPB Repeat Region of East African Leishmania donovani Strains. PLoS Negl Trop Dis 2013. DOI:10.1371/journal.pntd.0002031.

7 Gelanew T, Kuhls K, Hurissa Z, et al. Inference of population structure of leishmania donovani strains isolated from different ethiopian visceral leishmaniasis endemic areas. PLoS Negl Trop Dis 2010. DOI:10.1371/journal.pntd.0000889.

8 Gelanew T, Hailu A, Scho’nian G, Lewis MD, Miles MA, Yeo M. Multilocus sequence and microsatellite identification of intra-specific hybrids and ancestor-like donors among natural ethiopian isolates of leishmania donovani. Int J Parasitol 2014. DOI:10.1016/j.ijpara.2014.05.008.

9 Zackay A, Cotton JA, Sanders M, et al. Genome wide comparison of Ethiopian Leishmania donovani strains reveals differences potentially related to parasite survival. PLoS Genet 2018. DOI:10.1371/journal.pgen.1007133.

10 Chargui N, Amro A, Haouas N, et al. Population structure of Tunisian Leishmania infantum and evidence for the existence of hybrids and gene flow between genetically different populations. Int J Parasitol 2009. DOI:10.1016/j.ijpara.2008.11.016.

11 Ravel C, Cortes S, Pratlong F, Morio F, Dedet JP, Campino L. First report of genetic hybrids between two very divergent Leishmania species: Leishmania infantum and Leishmania major. Int J Parasitol 2006. DOI:10.1016/j.ijpara.2006.06.019.

12 Odiwuor S, De Doncker S, Maes I, Dujardin JC, Van der Auwera G. Natural Leishmania donovani/Leishmania aethiopica hybrids identified from Ethiopia. Infect Genet Evol 2011. DOI:10.1016/j.meegid.2011.04.026.

13 Ozbilgin A, Culha G, Zeyrek FY, et al. Cutaneous and visceral tropisms of Leishmania tropica/Leishmania infantum hybrids in a murine model: First report of hybrid Leishmania strains isolated in Turkey. Int J Infect Dis 2012. DOI:10.1016/j.ijid.2012.05.702.

14 Akopyants NS, Kimblin N, Secundino N, et al. Demonstration of genetic exchange during cyclical development of Leishmania in the sand fly vector. Science (80-) 2009. DOI: 10.1126/science.1169464.

15 Sadlova J, Yeo M, Seblova V, et al. Visualisation of leishmania donovani fluorescent hybrids during early stage development in the sand fly vector. PLoS One 2011. DOI: 10.1371/journal.pone.0019851.

16 Inbar E, Akopyants NS, Charmoy M, et al. The Mating Competence of Geographically Diverse Leishmania major Strains in Their Natural and Unnatural Sand Fly Vectors. PLoS Genet 2013. DOI:10.1371/journal.pgen.1003672.

17 Romano A, Inbar E, Debrabant A, et al. Cross-species genetic exchange between visceral and cutaneous strains of Leishmania in the sand fly vector. Proc Natl Acad Sci U S A 2014. DOI:10.1073/pnas.1415109111.

18 Inbar E, Shaik J, Iantorno SA, et al. Whole genome sequencing of experimental hybrids supports meiosis-like sexual recombination in leishmania. PLoS Genet 2019. DOI:10.1371/journal.pgen.1008042.

19 WHO. Status of endemicity of cutaneous leishmaniasis, worldwide, 2018. Geneva, 2018 https://www.who.int/leishmaniasis/burden/GHO_CL_2018.pdf?ua=1.

20 Karunaweera ND, Ginige S, Senanayake S, et al. Spatial Epidemiologic Trends and Hotspots of Leishmaniasis, Sri Lanka, 2001-2018. Emerg Infect Dis 2020; 26: 1–10.

21 Siriwardana HVYD, Karunanayake P, Goonerathne L, Karunaweera ND. Emergence of visceral leishmaniasis in Sri Lanka: a newly established health threat. Pathog Glob Health 2017. DOI:10.1080/20477724.2017.1361564.

22 WHO. Status of endemicity of visceral leishmaniasis, worldwide, 2018. Geneva, 2018 https://www.who.int/leishmaniasis/burden/GHO_VL_2018.pdf?ua=1.

23 Thakur L, Singh KK, Shanker V, et al. Atypical leishmaniasis: A global perspective with emphasis on the Indian subcontinent. PLoS Negl. Trop. Dis. 2018. DOI:10.1371/journal.pntd.0006659.

24 Lypaczewski P, Hoshizaki J, Zhang W-W, et al. A complete Leishmania donovani reference genome identifies novel genetic variations associated with virulence. Sci Rep 2018; 8. DOI:10.1038/s41598-018-34812-x.

25 Zhang WW, Ramasamy G, McCall LI, et al. Genetic Analysis of Leishmania donovani Tropism Using a Naturally Attenuated Cutaneous Strain. PLoS Pathog 2014; 10: e1004244.

26 Samarasinghe SR, Samaranayake N, Kariyawasam UL, Siriwardana YD, Imamura H, Karunaweera ND. Genomic insights into virulence mechanisms of Leishmania donovani: Evidence from an atypical strain. BMC Genomics 2018. DOI:10.1186/s12864-018-5271-z.

27 Leinonen R, Sugawara H, Shumway M. The sequence read archive. Nucleic Acids Res 2011. DOI:10.1093/nar/gkq1019.

28 Franssen SU, Durrant C, Stark O, et al. Global genome diversity of the Leishmania donovani complex. Elife 2020. DOI:10.7554/eLife.51243.

29 Imamura H, Downing T, van den Broeck F, et al. Evolutionary genomics of epidemic visceral leishmaniasis in the Indian subcontinent. Elife 2016. DOI:10.7554/eLife.12613.

30 Cuypers B, Berg M, Imamura H, et al. Integrated genomic and metabolomic profiling of ISC1, an emerging Leishmania donovani population in the Indian subcontinent. Infect Genet Evol 2018. DOI:10.1016/j.meegid.2018.04.021.

31 Cotton JA, Durrant C, Franssen SU, et al. Genomic analysis of natural intra-specific hybrids among Ethiopian isolates of Leishmania donovani. PLoS Negl Trop Dis 2020. DOI:10.1371/journal.pntd.0007143.

32 Koboldt DC, Zhang Q, Larson DE, et al. VarScan 2: Somatic mutation and copy number alteration discovery in cancer by exome sequencing. Genome Res 2012; 22: 568–76.

33 Aslett M, Aurrecoechea C, Berriman M, Al. E. TriTrypDB: a functional genomic resource for the Trypanosomatidae. Nucleic Acids Res 2010; 38: D457–62.

34 Camacho E, González-de la Fuente S, Rastrojo A, et al. Complete assembly of the Leishmania donovani (HU3 strain) genome and transcriptome annotation. Sci Rep 2019. DOI: 10.1038/s41598-019-42511-4.

35 Sterkers Y, Crobu L, Lachaud L, Pagès M, Bastien P. Parasexuality and mosaic aneuploidy in Leishmania: Alternative genetics. Trends Parasitol. 2014. DOI:10.1016/j.pt.2014.07.002.

36 Barja PP, Pescher P, Bussotti G, et al. Haplotype selection as an adaptive mechanism in the protozoan pathogen Leishmania donovani. Nat Ecol Evol 2017. DOI: 10.1038/s41559-017-0361-x.

37 Senanayake SASC, Abeyewicreme W, Dotson EM, Karunaweera ND. Characteristics of phlebotomine sandflies in selected areas of Sri Lanka. Southeast Asian J Trop Med Public Health 2015.

38 Volf P, Benkova I, Myskova J, Sadlova J, Campino L, Ravel C. Increased transmission potential of Leishmania major/Leishmania infantum hybrids. Int J Parasitol 2007. DOI:10.1016/j.ijpara.2007.02.002.

39 Lypaczewski P, Zhang W-W, Matlashewski G. A single amino acid mutation in the RagC protein of Leishmania donovani alters pathogenesis and tropism (In preparation). 2020.

40 Siriwardana Y, Deepachandi B, Weliange SDS, et al. First Evidence for Two Independent and Different Leishmaniasis Transmission Foci in Sri Lanka: Recent Introduction or Long-Term Existence? J Trop Med 2019. DOI:10.1155/2019/6475939.

41 Siriwardana Y, Zhou G, Deepachandi B, et al. Trends in Recently Emerged Leishmania donovani Induced Cutaneous Leishmaniasis, Sri Lanka, for the First 13 Years. Biomed Res Int 2019. DOI:10.1155/2019/4093603.

42 Mahmoudzadeh-Niknam H, Kiaei SS, Iravani D. Leishmania tropica infection, in comparison to Leishmania major, induces lower delayed type hypersensitivity in BALB/c mice. Korean J Parasitol 2007. DOI:10.3347/kjp.2007.45.2.103.

43 Croft SL, Sundar S, Fairlamb AH. Drug resistance in leishmaniasis. Clin. Microbiol. Rev. 2006. DOI:10.1128/CMR.19.1.111-126.2006.

44 Iantorno SA, Durrant C, Khan A, et al. Gene expression in Leishmania is regulated predominantly by gene dosage. MBio 2017. DOI:10.1128/mBio.01393-17.

45 Kassahun A, Sadlova J, Benda P, et al. Natural infection of bats with Leishmania in Ethiopia. Acta Trop 2015. DOI:10.1016/j.actatropica.2015.07.024.

46 Kassahun A, Sadlova J, Dvorak V, et al. Detection of leishmania donovani and L. tropica in ethiopian wild rodents. Acta Trop 2015. DOI: 10.1016/j.actatropica.2015.02.006.

47 Fernando SD, Dharmawardana P, Semege S, et al. The risk of imported malaria in security forces personnel returning from overseas missions in the context of prevention of re-introduction of malaria to Sri Lanka. Malar J 2016. DOI:10.1186/s12936-016-1204-y.

48 Li H. Aligning sequence reads, clone sequences and assembly contigs with BWA-MEM. 2013; arXiv:1303.

49 Li H, Handsaker B, Wysoker A, et al. The Sequence Alignment/Map format and SAMtools. Bioinformatics 2009; 25: 2078–9.

50 Bradbury PJ, Zhang Z, Kroon DE, Casstevens TM, Ramdoss Y, Buckler ES. TASSEL: Software for association mapping of complex traits in diverse samples. Bioinformatics 2007. DOI: 10.1093/bioinformatics/btm308.

51 Letunic I, Bork P. Interactive Tree Of Life (iTOL) v4: recent updates and new developments. Nucleic Acids Res 2019. DOI:10.1093/nar/gkz239.

52 Danecek P, McCarthy SA. BCFtools/csq: Haplotype-aware variant consequences. Bioinformatics 2017. DOI: 10.1093/bioinformatics/btx100.

53 Krzywinski M, Schein J, Birol I, et al. Circos: An information aesthetic for comparative genomics. Genome Res 2009. DOI:10.1101/gr.092759.109.

54 EMBL-EBI H. Clustal Omega: Multiple Sequence Alignment. Eur Mol Biol Lab 2017.

