## Supplementary Information for "Evidence that interspecies *Leishmania* hybrids contribute to changes in disease pathology"

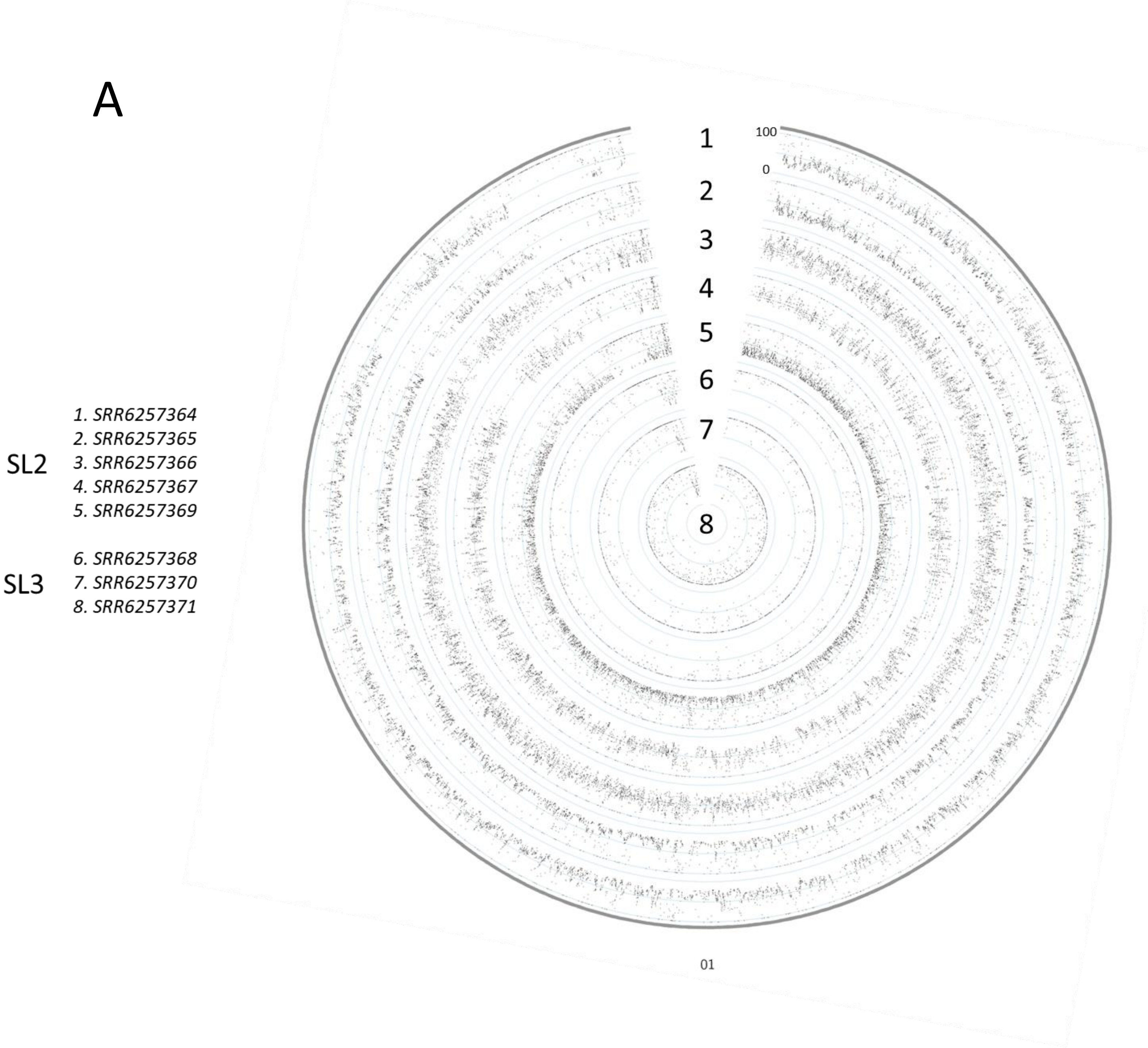

B

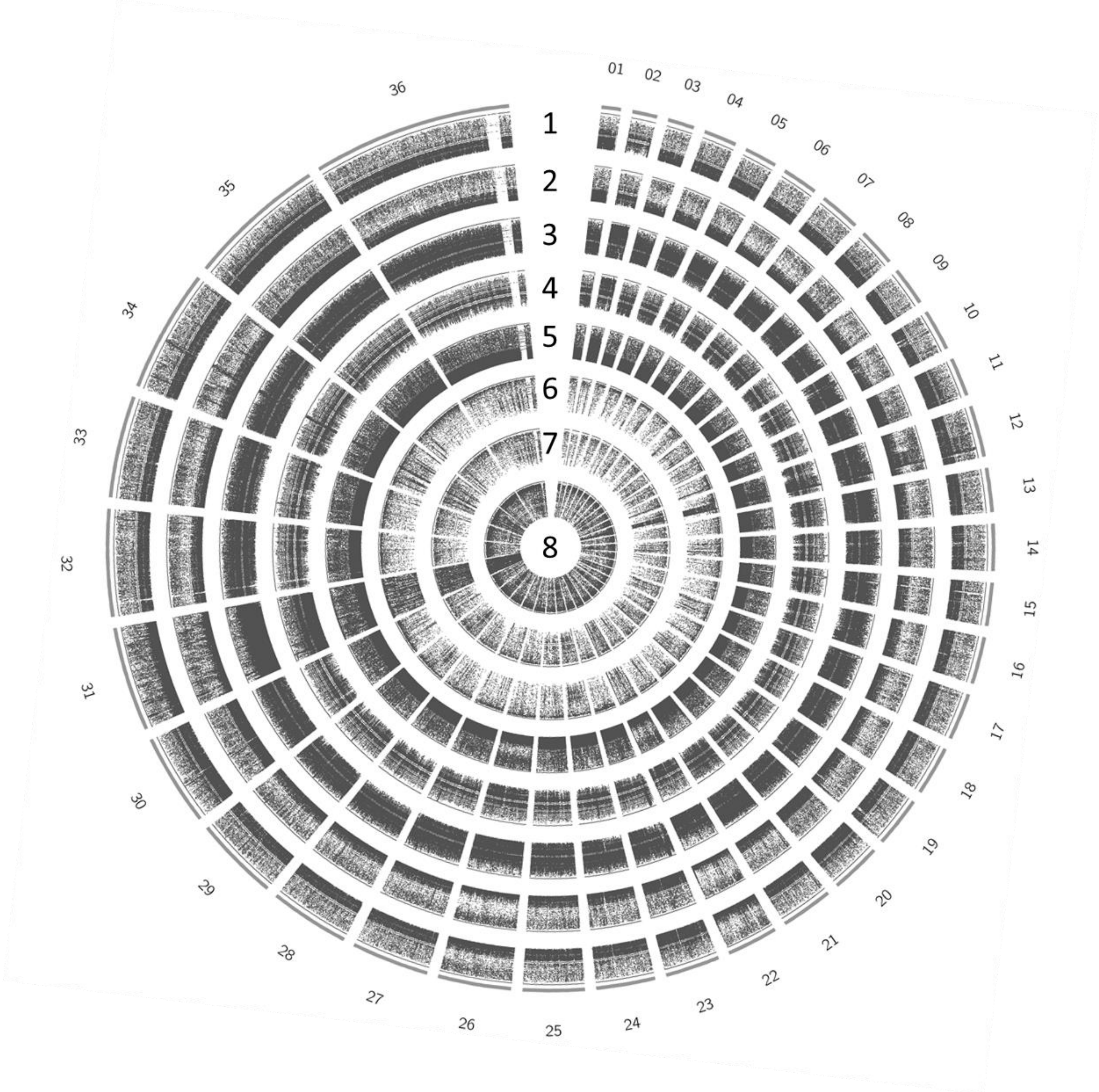


*1. SRR6257364*
*2. SRR6257365*
*3. SRR6257366*
*4. SRR6257367*
*5. SRR6257369*
*6. SRR6257368*
*7. SRR6257370*
*8. SRR6257371*

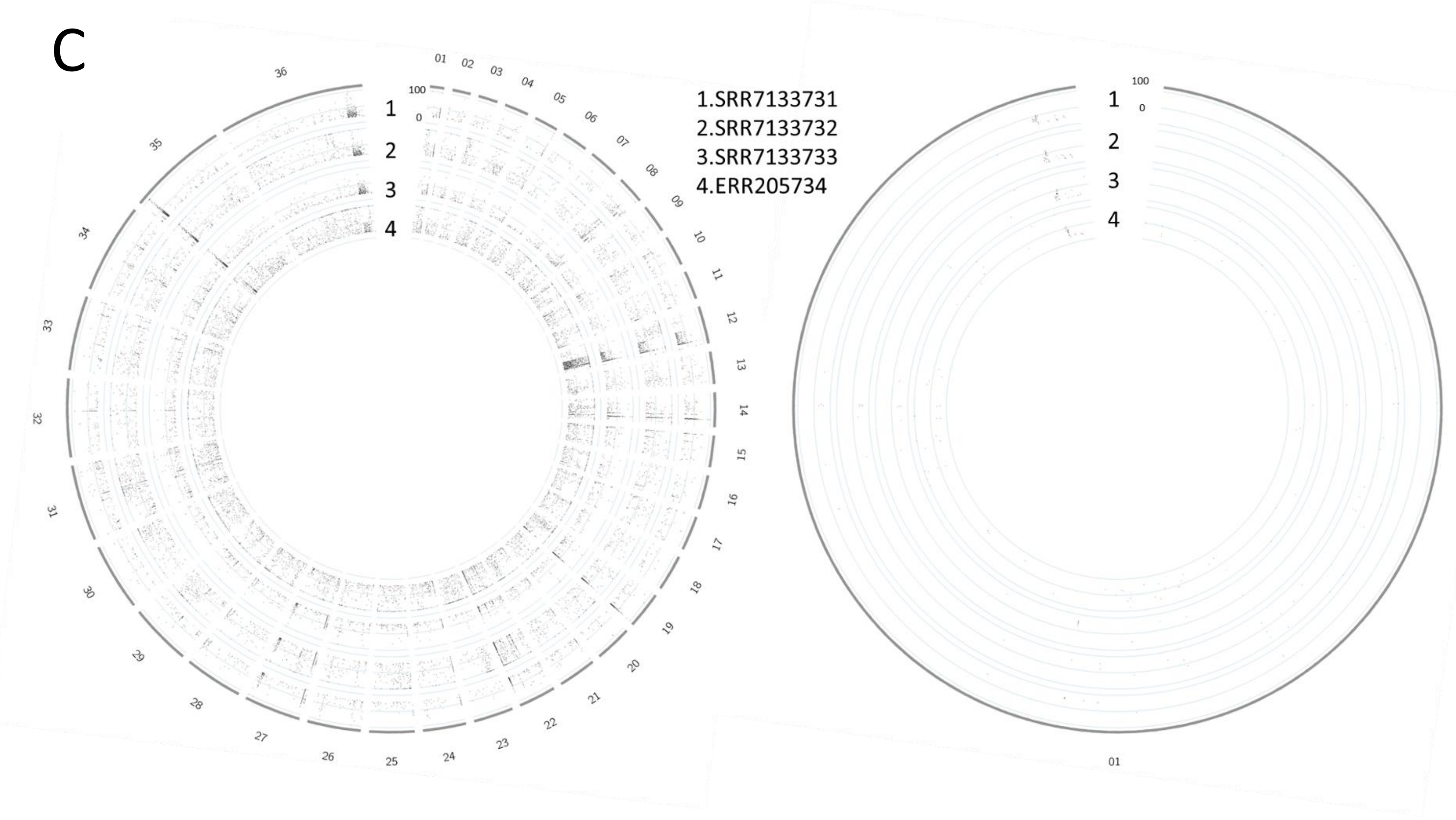


**Supplementary Figure S1.** Frequency and distribution of polymorphisms Sri Lankan isolates. The location of every polymorphism is depicted along the X-axis and the underlying read frequency along the Y-axis as a single dot. Points centered around the center of each track represent ~50%, points at the outer edge of each track represent homozygous ~100% polymorphisms. To better distinguish the frequency distribution, the plot is limited to Chromosome 1 (A) and observe the distribution of polymorphism across the genome, all 36 chromosomes (B) in the SL2 and SL3 groups. **C.** Location and read frequencies of SL1 group isolates across the whole genome or Chromosome 1.

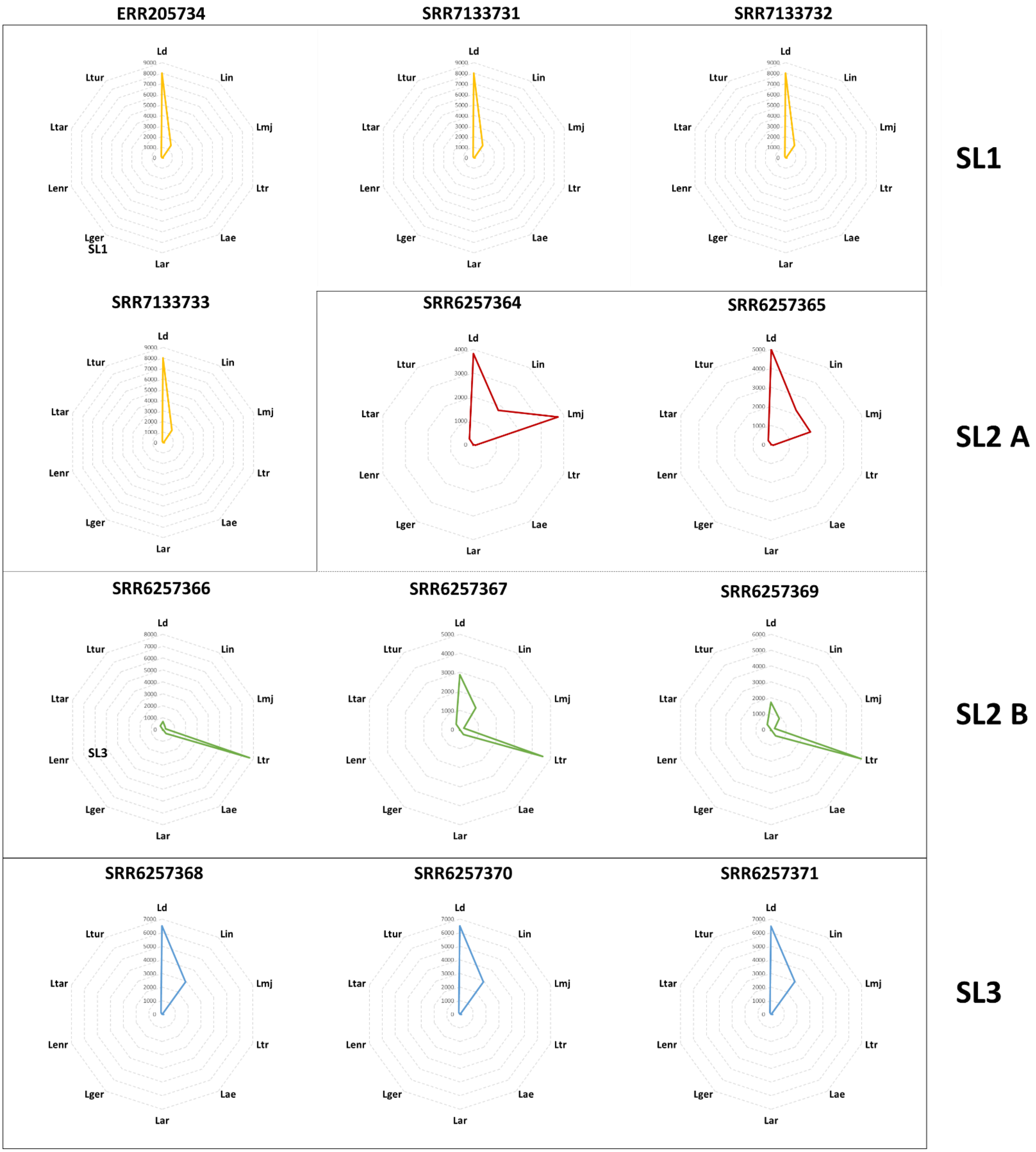


**Supplementary Figure S2.** Distribution of species origin of reconstructed genes as determined by BLAST analysis.

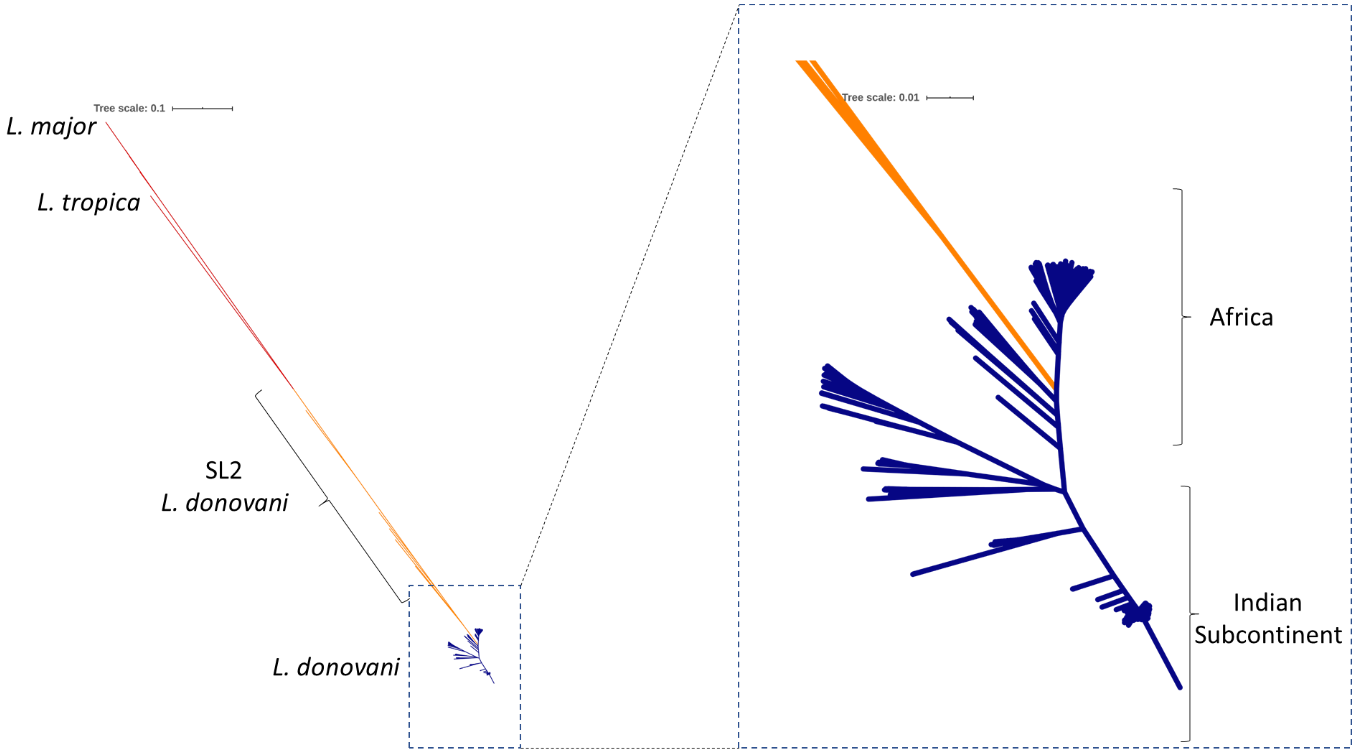


**Supplementary Figure S3.** Phylogenetic analysis of *L. donovani* isolates, *L. major* and *L. tropica.* Branches corresponding to *L. donovani* are colored in blue, *L. major* and *L. tropica* in red, and SL2 putative hybrid parasites in orange.

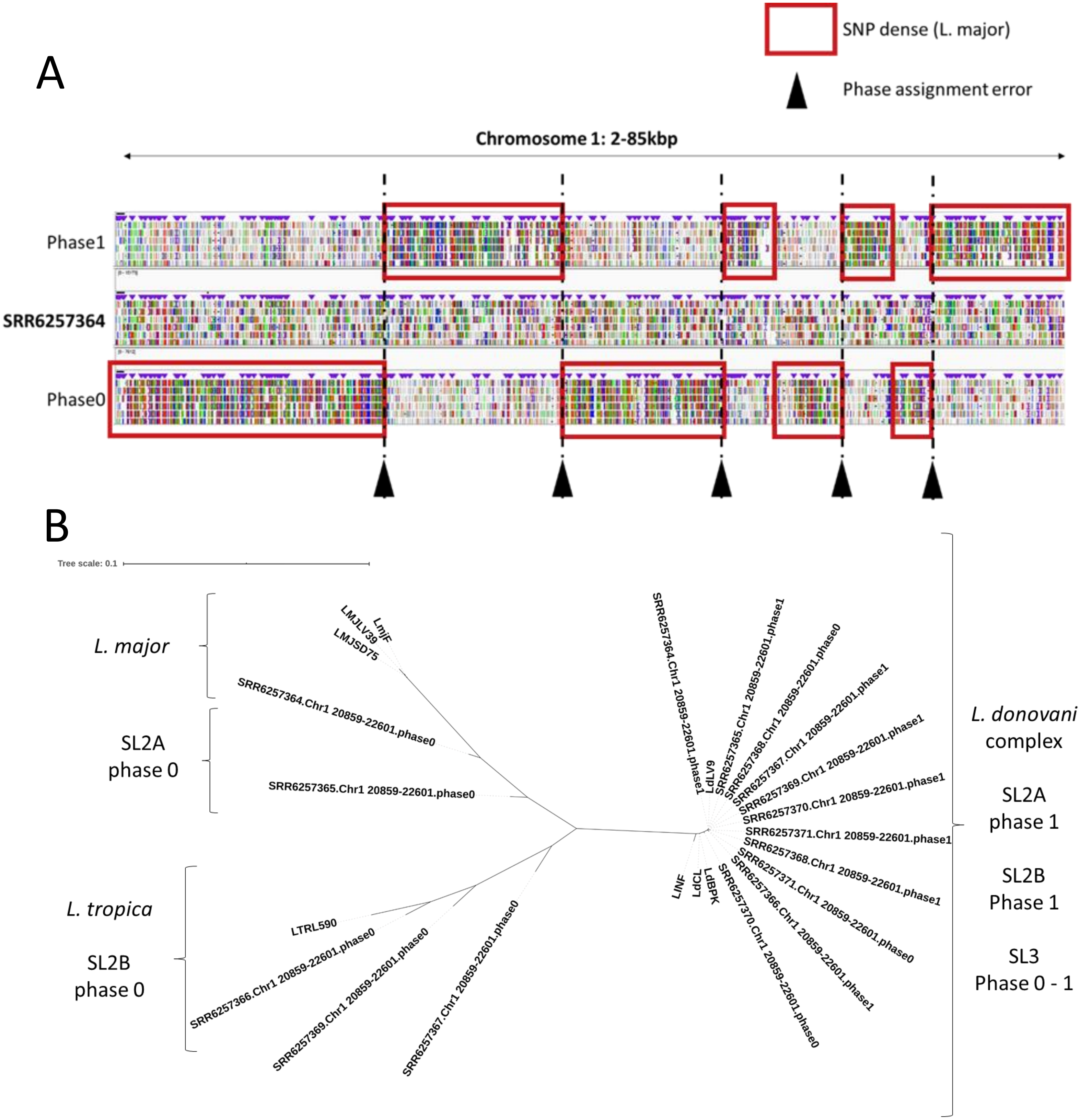


**Supplementary Figure S4.** Phylogenetic analysis after read-based phasing to separate haplotypes. **A.** Read based phasing results in chimeric haplotypes as read blocks are assigned to the wrong phase sets as shown by the alternating assignment of SNP dense (*L. major*) read blocks in continuous phase sets. **B.** Phylogenetic analysis using phased haplotypes for a section of chromosome 1 showing a different clustering across phases in SL2A *L. major* hybrids and SL2B *L. tropica* hybrids.

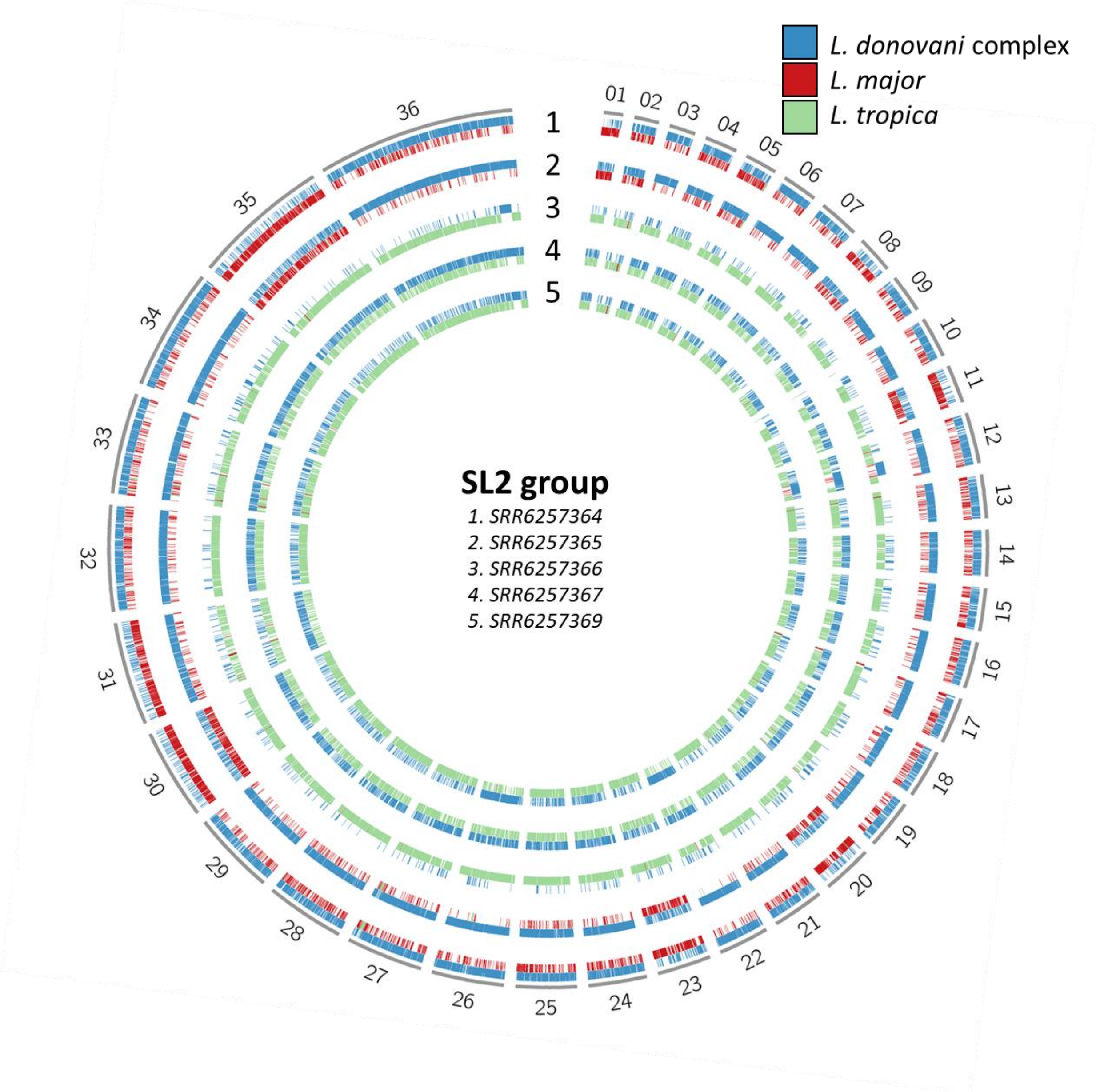


**Supplementary Figure S5.** Representation of gene ancestry across the entire genome. Genes of *L. donovani* species complex origin are marked in blue. Genes with hybrid ancestry (*L. major & L. donovani*, or *L. tropica & L. donovani*) are colored in red and green respectively.

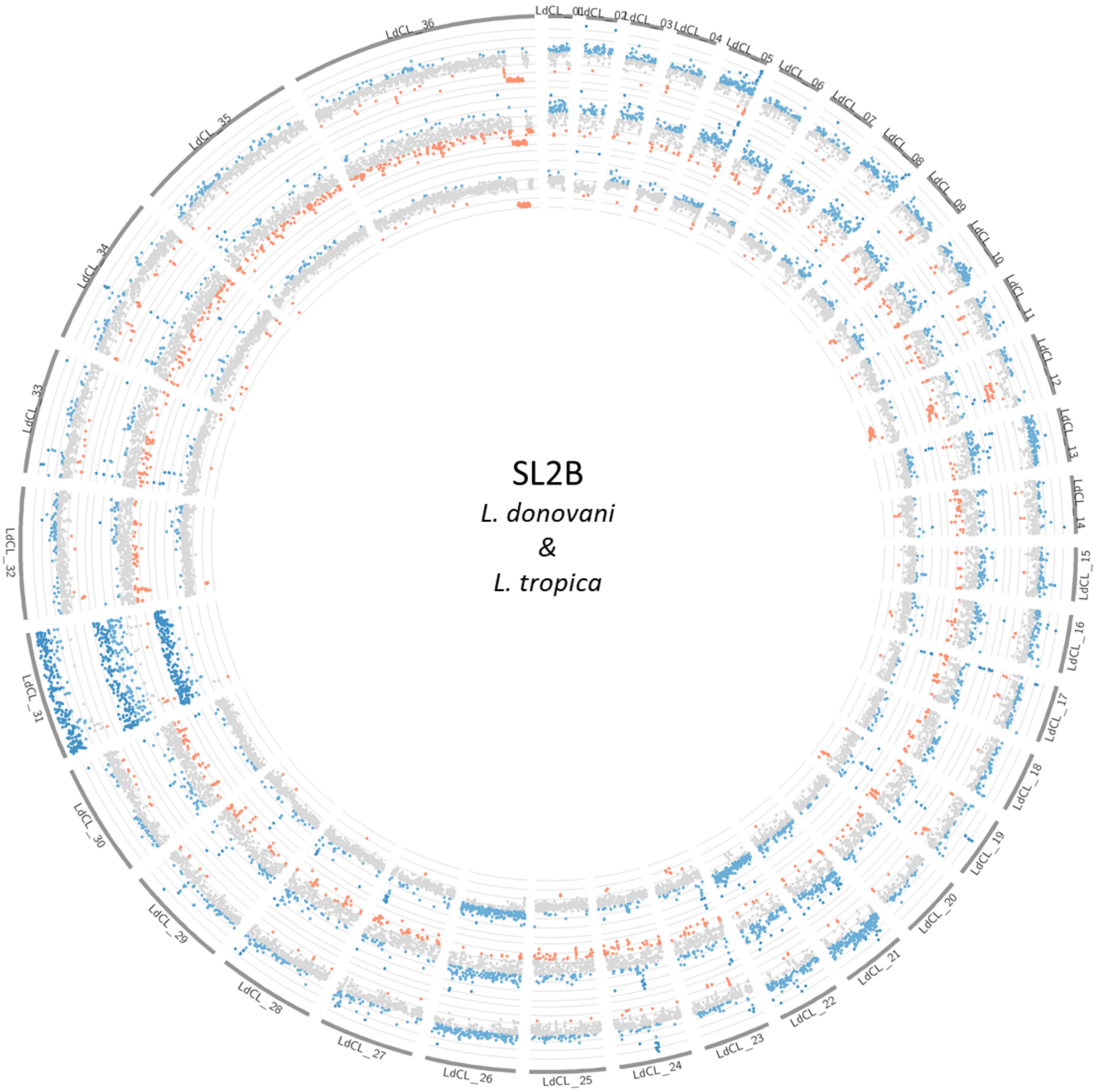


**Supplementary Figure S5.** Comparison of chromosomal aneuploidy in SL2B (*L. tropica* hybrids) isolates.

**Supplementary Table S1.** List of accession numbers used in this study

| **SRA Accession Numbers** |
|---|
| ERR018830 ERR018831 ERR018832 ERR018833 ERR018834 ERR018835 ERR018836 |
| ERR018837 ERR018838 ERR018839 ERR018840 ERR018841 ERR018842 ERR018843 |
| ERR018844 ERR018845 ERR018846 ERR018847 ERR018849 ERR018850 ERR018851 |
| ERR018852 ERR018853 ERR018854 ERR018855 ERR018856 ERR018857 ERR018859 |
| ERR018860 ERR018861 ERR018862 ERR019524 ERR034092 ERR036057 ERR036058 |
| ERR1329883 ERR1329884 ERR205723 ERR205724 ERR205726 ERR205729 ERR205730 |
| ERR205731 ERR205732 ERR205733 ERR205734 ERR205735 ERR205736 ERR205737 |
| ERR205738 ERR205739 ERR205740 ERR205741 ERR205742 ERR205743 ERR205744 |
| ERR205745 ERR205746 ERR205747 ERR205748 ERR205749 ERR205750 ERR205751 |
| ERR205752 ERR205753 ERR205754 ERR205755 ERR205756 ERR205757 ERR205758 |
| ERR205759 ERR205760 ERR205761 ERR205763 ERR205765 ERR205766 ERR205767 |
| ERR205768 ERR205770 ERR205771 ERR205772 ERR205774 ERR205775 ERR205776 |
| ERR205777 ERR205778 ERR205779 ERR205780 ERR205781 ERR205782 ERR205783 |
| ERR205784 ERR205785 ERR205786 ERR205787 ERR205788 ERR205789 ERR205790 |
| ERR205791 ERR205792 ERR205793 ERR205794 ERR205795 ERR205797 ERR205799 |
| ERR205800 ERR205801 ERR205802 ERR205803 ERR205804 ERR205805 ERR205806 |
| ERR205807 ERR205808 ERR205809 ERR205810 ERR205811 ERR205812 ERR205813 |
| ERR205814 ERR205815 ERR205816 ERR205817 ERR205818 ERR205819 ERR205820 |
| ERR206260 ERR206261 ERR206262 ERR206263 ERR206264 ERR206265 ERR206266 |
| ERR206267 ERR206268 ERR206269 ERR206270 ERR206271 ERR206272 ERR206273 |
| ERR206274 ERR206275 ERR206276 ERR206277 ERR206278 ERR206279 ERR206280 |
| ERR206281 ERR206282 ERR206283 ERR206284 ERR206285 ERR206286 ERR206287 |
| ERR206288 ERR206289 ERR206290 ERR206291 ERR206292 ERR206293 ERR206294 |
| ERR206295 ERR206296 ERR206297 ERR206298 ERR206299 ERR206300 ERR206301 |
| ERR206302 ERR206303 ERR206304 ERR206305 ERR206306 ERR206307 ERR206308 |
| ERR206309 ERR206310 ERR206311 ERR206312 ERR206313 ERR206314 ERR206315 |
| ERR206316 ERR206317 ERR206318 ERR206319 ERR206320 ERR206321 ERR206322 |
| ERR206323 ERR206324 ERR206325 ERR206326 ERR206327 ERR206328 ERR206329 |
| ERR206330 ERR206331 ERR206332 ERR206333 ERR206334 ERR206335 ERR206336 |
| ERR206337 ERR206338 ERR206339 ERR206340 ERR206341 ERR206342 ERR206343 |
| ERR206344 ERR206345 ERR206346 ERR206347 ERR206348 ERR206349 ERR206350 |
| ERR206351 ERR206352 ERR206353 ERR206354 ERR206355 ERR206356 ERR206357 |
| ERR206358 ERR206359 ERR206360 ERR206361 ERR206362 ERR206363 ERR206364 |
| ERR206365 ERR206366 ERR206367 ERR206368 ERR206369 ERR206370 ERR206371 |
| ERR206372 ERR206373 ERR206374 ERR206375 ERR206376 ERR206377 ERR206378 |
| ERR206379 ERR206380 ERR206381 ERR206382 ERR206383 ERR206384 ERR206385 |
| ERR206386 ERR206387 ERR206388 ERR206389 ERR206390 ERR206391 ERR206392 |
| ERR206393 ERR206394 ERR206395 ERR206396 ERR206397 ERR206398 ERR206399 |
| ERR206400 ERR206401 ERR206402 ERR206403 ERR206404 ERR206405 ERR206406 |
| ERR206407 ERR206408 ERR206409 ERR206410 ERR206411 ERR206412 ERR206413 |
| ERR206414 ERR206415 ERR206416 ERR206417 ERR206418 ERR206419 ERR206420 |
| ERR206421 ERR206422 ERR206423 ERR206424 ERR206425 ERR206426 ERR206427 |
| ERR206428 ERR206429 ERR206430 ERR206431 ERR206432 ERR206433 ERR206434 |
| ERR206435 ERR206436 ERR206437 ERR206438 ERR206439 ERR206440 ERR206441 |
| ERR206442 ERR206443 ERR206444 ERR206445 ERR206446 ERR206447 ERR206448 |
| ERR206449 ERR206450 ERR206451 ERR206452 ERR206453 ERR206454 ERR206455 |
| ERR206456 ERR206457 ERR206458 ERR206459 ERR206460 ERR206461 ERR206462 |

ERR206463 ERR206464 ERR206465 ERR206466 ERR206467 ERR206468 ERR206469
ERR206470 ERR206471 ERR206472 ERR206473 ERR206474 ERR206475 ERR206476
ERR206477 ERR206478 ERR206479 ERR206480 ERR206481 ERR206482 ERR206483
ERR206484 ERR206485 ERR206486 ERR206487 ERR206488 ERR206489 ERR206490
ERR206491 ERR206492 ERR206493 ERR206494 ERR206495 ERR206496 ERR206497
ERR206498 ERR206499 ERR206500 ERR206501 ERR206502 ERR206503 ERR206504
ERR206505 ERR206506 ERR206507 ERR206508 ERR206509 ERR206510 ERR206511
ERR206512 ERR206513 ERR206514 ERR206515 ERR206516 ERR206517 ERR206518
ERR206519 ERR206520 ERR206521 ERR206522 ERR206523 ERR206524 ERR206525
ERR206526 ERR206527 ERR206528 ERR206529 ERR206530 ERR206531 ERR206532
ERR206533 ERR206534 ERR206535 ERR206536 ERR206537 ERR206538 ERR206539
ERR206540 ERR206541 ERR206542 ERR206543 ERR206544 ERR206545 ERR206546
ERR206547 ERR206548 ERR206549 ERR206550 ERR206551 ERR206552 ERR206553
ERR206554 ERR206555 ERR206556 ERR206557 ERR206558 ERR206559 ERR206560
ERR206561 ERR206562 ERR206563 ERR206564 ERR206565 ERR206566 ERR206567
ERR206568 ERR206569 ERR206570 ERR206571 ERR206573 ERR206574 ERR206575
ERR206576 ERR206577 ERR206578 ERR206579 ERR206580 ERR206581 ERR206582
ERR206583 ERR206584 ERR206585 ERR206586 ERR206587 ERR206588 ERR206589
ERR206590 ERR206591 ERR206592 ERR206593 ERR206594 ERR206595 ERR206597
ERR206598 ERR206599 ERR206600 ERR206601 ERR206602 ERR206603 ERR206604
ERR206605 ERR206606 ERR206607 ERR206608 ERR206609 ERR206610 ERR206611
ERR206612 ERR206613 ERR206614 ERR206615 ERR206616 ERR206617 ERR206618
ERR207770 ERR207771 ERR207772 ERR207773 ERR207774 ERR207775 ERR207776
ERR207777 ERR207778 ERR207779 ERR207780 ERR2124117 ERR2124119 ERR2124120
ERR2124121 ERR2124122 ERR2124123 ERR2124124 ERR2124125 ERR2124126 ERR2124127
ERR2124128 ERR261858 ERR261859 ERR261860 ERR261861 ERR261862 ERR261863
ERR261864 ERR261865 ERR261866 ERR261867 ERR261868 ERR304754 ERR304755
ERR304756 ERR304757 ERR304758 ERR304759 ERR304760 ERR304761 ERR304762
ERR304763 ERR304764 ERR304765 ERR304766 ERR305913 ERR305914 ERR305915
ERR305916 ERR305917 ERR310823 ERR310824 ERR310825 ERR310826 ERR310827
ERR310828 ERR310829 ERR310830 ERR310831 ERR310832 ERR310833 ERR310834
ERR310835 ERR310836 ERR310837 ERR363217 ERR363218 ERR363219 ERR363220
ERR363221 ERR363222 ERR363223 ERR363224 ERR363225 ERR363226 ERR363227
ERR363228 ERR363229 ERR363230 ERR363231 ERR363232 ERR363233 ERR363234
ERR363235 ERR363236 ERR3723986 ERR3723987 ERR3723988 ERR3723989 ERR376170
ERR376171 ERR376172 ERR376173 ERR376174 ERR376175 ERR376176 ERR376177
ERR376178 ERR376179 ERR376180 ERR376181 ERR376182 ERR376183 ERR376184
ERR376185 ERR376186 ERR412008 ERR438786 ERR438787 ERR438788 ERR438789
ERR438790 ERR438791 ERR438792 ERR438793 ERR438794 ERR438795 ERR438796
ERR539726 ERR539727 ERR539728 ERR539729 ERR539730 ERR539732 ERR539733
ERR539734 ERR539735 ERR539736 ERR539737 ERR539738 ERR539739 ERR541269
ERR541270 ERR541271 ERR541272 ERR541273 ERR541274 ERR541275 ERR541276
ERR568417 ERR568418 ERR568419 ERR600042 ERR600043 ERR600044 ERR600045
ERR600046 ERR600047 ERR600048 ERR600049 ERR600050 ERR600051 ERR600054
ERR600056 ERR754983 ERR754984 ERR862590 ERR862591 ERR862592 ERR862593
ERR862594 ERR862595 ERR862596 ERR862597 ERR925016 ERR925017 ERR925018
ERR925019 ERR925020 ERR925021 ERR925022 ERR925023 ERR925024 ERR925025
ERR925026 SRR1254938 SRR1254940 SRR6257364 SRR6257365 SRR6257366 SRR6257367
SRR6257368 SRR6257369 SRR6257370 SRR6257371 SRR6316153 SRR6316158 SRR6316159
SRR6316161 SRR6369639 SRR6369648 SRR6369649 SRR6369650 SRR6369651 SRR6369652
SRR6369653 SRR6369660 SRR7133731 SRR7133732 SRR7133733

**Supplementary Table S2.** Summary of polymorphisms across Sri Lankan isolates

| Group | Isolate | Homozygous | Heterozygous |
|---|---|---|---|
| **SL1*** | ERR205734 | 34401 | 13075 |
| | SRR7133731 | 20043 | 4987 |
| | SRR7133732 | 17732 | 4754 |
| | SRR7133733 | 38481 | 10187 |
| **SL2** | SRR6257364 | 96335 | 824077 |
| | SRR6257365 | 128317 | 634899 |
| | SRR6257366 | 268534 | 1559016 |
| | SRR6257367 | 163181 | 790134 |
| | SRR6257369 | 110673 | 1083904 |
| **SL3** | SRR6257368 | 143212 | 33995 |
| | SRR6257370 | 141234 | 28620 |
| | SRR6257371 | 141145 | 33266 |

*: SL1 group isolates were compared to the LdBPK reference genome to avoid masking SL1 specific homozygous polymorphism as described in Methods

**Supplementary Table S3. Non-synonymous polymorphisms identified in the SL3 group in common with *L. major* SNPs**

| Chr | Position | Gene ID | Protein Name | Effect |
|---|---|---|---|---|
| LdCL_06 | 373712 | LdCL_060013900 | hypothetical protein | Ser2073Thr |
| LdCL_06 | 373865 | | | Ala2124Ser |
| LdCL_06 | 374067 | | | Thr2191Met |
| LdCL_06 | 374691 | | | Leu2399Ser |
| LdCL_06 | 375785 | | | Leu2764Met |
| LdCL_06 | 379939 | LdCL_060014000 | dihydrofolate reductase-thymidylate synthase | Thr200Ala |
| LdCL_06 | 379942 | | | Thr201Ala |
| LdCL_06 | 382826 | LdCL_060014100 | arginine N-methyltransferase, type III, putative | Glu64Gly |
| LdCL_06 | 401369 | LdCL_060014600 | hypothetical protein | Cys267Tyr |
| LdCL_06 | 449436 | LdCL_060016100 | 2OG-Fe(II) oxygenase superfamily, putative | Ala8Val |
| LdCL_06 | 483218 | LdCL_060017300 | protein kinase, putative | His378Arg |
| LdCL_07 | 362713 | LdCL_070013400 | hypothetical protein | Asp33Gly |
| LdCL_09 | 135099 | LdCL_090009000 | DNA photolyase, putative | Arg131Gln |
| LdCL_09 | 163075 | LdCL_090009800 | tyrosine phosphatase, putative | Ala771_Ala774del |
| LdCL_09 | 275934 | LdCL_090013200 | hypothetical protein | Lys504Glu |
| LdCL_10 | 169505 | LdCL_100009100 | pteridine transporter, putative | Thr174Arg |
| LdCL_11 | 401388 | LdCL_110016000 | hypothetical protein | Gln72Pro |
| LdCL_11 | 401414 | | | His81Tyr |
| LdCL_11 | 490683 | LdCL_110018100 | ATP-binding cassette subfamily A, member 1, putative | Gly631Asp |
| LdCL_12 | 431663 | LdCL_120013900 | hypothetical protein | Tyr43His |
| LdCL_12 | 569653 | LdCL_120017400 | hypothetical protein | Tyr43His |
| LdCL_12 | 569675 | | | Trp50* |
| LdCL_12 | 588162 | LdCL_120017900 | surface antigen protein 2, putative | Gly101Ser |
| LdCL_12 | 588178 | | | Thr106Asn |
| LdCL_12 | 589089 | | | Asp410Asn |
| LdCL_12 | 589891 | | | Ala677Val |
| LdCL_12 | 605397 | LdCL_120018400 | surface antigen protein 2, putative | Glu76Val |
| LdCL_12 | 605472 | | | Asp101Gly |
| LdCL_12 | 605487 | | | Thr106Asn |
| LdCL_12 | 605492 | | | Met108Val |
| LdCL_12 | 605503 | | | His111Gln |
| LdCL_12 | 605535 | | | Ser122Ile |
| LdCL_12 | 605550 | | | Ala127Asp |
| LdCL_12 | 605563 | | | Asn131Lys |
| LdCL_12 | 605577 | | | Ser136Thr |

| | | | | |
|---|---|---|---|---|
| LdCL_12 | 605580 | | | Ser137Leu |
| LdCL_12 | 605582 | | | Val138Leu |
| LdCL_13 | 374514 | LdCL_130015300 | leucyl-tRNA synthetase, putative | Leu308Phe |
| LdCL_14 | 222212 | LdCL_140011500 | hypothetical protein | Thr687Ala |
| LdCL_14 | 346848 | LdCL_140014000 | hypothetical protein | Gln2642dup |
| LdCL_14 | 534346 | LdCL_140017800 | kinesin K39, putative | Ser1303Asn |
| LdCL_14 | 538404 | LdCL_140017800 | kinesin K39, putative | Ala2656Thr |
| LdCL_14 | 590582 | LdCL_140019500 | NADP oxidoreductase coenzyme F420-dependent, putative | Val47Ala |
| LdCL_14 | 606148 | LdCL_140019900 | serine hydroxymethyltransferase (SHMT-L) | Ser24Gly |
| LdCL_14 | 659681 | LdCL_140021300 | hypothetical protein | Ala747Val |
| LdCL_15 | 136121 | LdCL_150008900 | hypothetical protein | Asn394Ser |
| LdCL_15 | 162995 | LdCL_150010100 | tb-292 membrane associated protein-like protein | Glu510Gln |
| LdCL_15 | 164150 | LdCL_150010100 | tb-292 membrane associated protein-like protein | Glu895Gln |
| LdCL_16 | 196816 | LdCL_160010600 | AAA domain containing protein, putative | Ser85Pro |
| LdCL_17 | 88351 | LdCL_170007500 | receptor-type adenylate cyclase, putative | Ser20Pro |
| LdCL_17 | 112477 | LdCL_170008200 | Kinesin motor domain containing protein, putative | Arg524Cys |
| LdCL_17 | 118974 | LdCL_170008300 | WD domain, G-beta repeat, putative | Ala399Val |
| LdCL_17 | 119553 | LdCL_170008300 | WD domain, G-beta repeat, putative | Phe206Ser |
| LdCL_18 | 609144 | LdCL_180019300 | pumilio protein 2, putative | Asn875Thr |
| LdCL_18 | 609173 | LdCL_180019300 | pumilio protein 2, putative | Ser885Gly |
| LdCL_18 | 609201 | LdCL_180019300 | pumilio protein 2, putative | Thr894Met |
| LdCL_19 | 478877 | LdCL_190016300 | Ankyrin repeats (3 copies)/Ankyrin repeat, putative | Gly3850Asp |
| LdCL_19 | 480896 | LdCL_190016300 | Ankyrin repeats (3 copies)/Ankyrin repeat, putative | Leu4525del |
| LdCL_22 | 551683 | LdCL_220018200 | hypothetical protein | Asn183Ser |
| LdCL_23 | 467160 | LdCL_230018000 | hypothetical protein | Val2766Ala |
| LdCL_24 | 154090 | LdCL_240009300 | Protein of unknown function (DUF3184), putative | Ser362Ala |
| LdCL_24 | 161257 | LdCL_240009400 | Protein of unknown function (DUF3184), putative | Ser357Ala |
| LdCL_24 | 324959 | LdCL_240014200 | hypothetical protein | Ser600Pro |
| LdCL_24 | 512669 | LdCL_240019700 | hypothetical protein | His426Asp |
| LdCL_24 | 734342 | LdCL_240025400 | WD domain, G-beta repeat/PFU (PLAA family ubiquitin binding), putative | Ser402Gly |
| LdCL_26 | 800281 | LdCL_260026400 | hypothetical protein | Ala209Thr |
| LdCL_26 | 884803 | LdCL_260028200 | kynureninase, putative | Ala189Val |
| LdCL_27 | 318494 | LdCL_270012900 | calcium uniporter protein, | Arg37His |

| | | | | |
|---|---|---|---|---|
| | | | mitochondrial, putative | |
| LdCL_27 | 321591 | LdCL_270013000 | MatE, putative | Ser8Gly |
| LdCL_27 | 321790 | | | Arg74His |
| LdCL_27 | 334538 | LdCL_270013300 | Right handed beta helix region/Periplasmic copper-binding protein (NosD), putative | Gln1101del |
| LdCL_27 | 581562 | LdCL_270019400 | protein kinase, putative | Ser1202Pro |
| LdCL_27 | 701974 | LdCL_270022700 | molybdopterin synthase sulfurylase-like protein, putative | Val389Ala |
| LdCL_27 | 893453 | LdCL_270026900 | Transmembrane adaptor Erv26, putative | Arg179Gln |
| LdCL_28 | 711944 | LdCL_280024500 | DNA repair protein-like protein | Asp652Gly |
| LdCL_28 | 890779 | LdCL_280029500 | mitochondrial DNA topoisomerase II | Pro1380Ser |
| LdCL_28 | 938512 | LdCL_280030800 | tRNA methyltransferase complex GCD14 subunit/Protein-L-isoaspartate(D-aspartate) O-methyltransfer... | Thr43Ile |
| LdCL_29 | 291220 | LdCL_290013000 | hypothetical protein | Arg423Leu |
| LdCL_29 | 496404 | LdCL_290018500 | hypothetical protein | Ala883Val |
| LdCL_29 | 677115 | LdCL_290021200 | hypothetical protein | Gly296Ser |
| LdCL_29 | 775470 | LdCL_290023400 | ATP-binding cassette protein subfamily H, member 2, putative | Leu183Ile |
| LdCL_29 | 1169010 | LdCL_290033900 | serine/threonine-protein kinase Nek, putative | Pro364Ser |
| LdCL_30 | 516744 | LdCL_300020600 | p1/s1 nuclease | Ala128Ser |
| LdCL_30 | 711383 | LdCL_300025400 | hypothetical protein | Leu122Arg |
| LdCL_31 | 168328 | LdCL_310010100 | amastin, putative | Ile78Val |
| LdCL_31 | 253938 | LdCL_310012600 | hypothetical protein | Tyr101His |
| LdCL_31 | 375755 | LdCL_310015700 | sodium stibogluconate resistance protein, putative | Glu458Gly |
| LdCL_31 | 449919 | LdCL_310017200 | hypothetical protein | Ala513Val |
| LdCL_31 | 466862 | LdCL_310017700 | hypothetical protein | Leu91Phe |
| LdCL_31 | 528422 | LdCL_310019500 | hypothetical protein | Leu1091Phe |
| LdCL_31 | 613221 | LdCL_310020500 | Cold-shock DNA-binding domain containing protein, putative | Leu31Pro |
| LdCL_31 | 619290 | LdCL_310020600 | hypothetical protein | Asn338Asp |
| LdCL_31 | 619307 | | | Asn332Ser |
| LdCL_31 | 619502 | | | Arg267Lys |
| LdCL_31 | 619821 | | | Thr161Ala |
| LdCL_31 | 619989 | | | Met105Val |
| LdCL_31 | 620189 | | | Thr38Met |
| LdCL_31 | 917260 | LdCL_310026100 | protein kinase, putative | Lys33Gln |

| | | | | |
|---|---|---|---|---|
| LdCL_31 | 987807 | LdCL_310027400 | RING-variant domain containing protein, putative | Ser104Asn |
| LdCL_31 | 987835 | LdCL_310027400 | RING-variant domain containing protein, putative | Leu95Phe |
| LdCL_31 | 1084057 | LdCL_310029900 | Fungal tRNA ligase phosphodiesterase domain containing protein, putative | Arg520Gln |
| LdCL_31 | 1151019 | LdCL_310030900 | hypothetical protein | Asn1902Ser |
| LdCL_31 | 1158599 | LdCL_310031000 | hypothetical protein | Asp359Asn |
| LdCL_31 | 1265045 | LdCL_310033400 | lipase, putative | Met346Ile |
| LdCL_31 | 1478843 | LdCL_310040100 | iron/zinc transporter protein-like protein | Cys269Tyr |
| LdCL_31 | 1482804 | LdCL_310040200 | iron/zinc transporter protein-like protein | Cys269Tyr |
| LdCL_32 | 221657 | LdCL_320011500 | ATP-dependent DEAD/H RNA helicase, putative | Gly19Glu |
| LdCL_32 | 844420 | LdCL_320027800 | hypothetical protein | Arg135His |
| LdCL_32 | 1135510 | LdCL_320035800 | Prefoldin subunit, putative | Thr370Met |
| LdCL_33 | 250875 | LdCL_330013100 | POT family, putative | Ser215Thr |
| LdCL_33 | 345811 | LdCL_330015500 | dnaj chaperone-like protein | Ala533Val |
| LdCL_33 | 350414 | LdCL_330015800 | hypothetical protein | Lys148Arg |
| LdCL_33 | 354193 | LdCL_330015900 | hypothetical protein | Thr59Ser |
| LdCL_33 | 562350 | LdCL_330021500 | protein kinase, putative | Cys1720Tyr |
| LdCL_34 | 168645 | LdCL_340009900 | hypothetical protein | Leu118Phe |
| LdCL_34 | 209343 | LdCL_340010500 | Amastin surface glycoprotein, putative | Ala41Gly |
| LdCL_34 | 226078 | LdCL_340010900 | phosphoglycan beta 1,2 arabinosyltransferase, (SCA like) | Thr81Ile |
| LdCL_34 | 372801 | LdCL_340013900 | serine/threonine-protein phosphatase PP1, putative | His99Gln |
| LdCL_34 | 507010 | LdCL_340017500 | amastin-like surface protein, putative | Leu15Phe |
| LdCL_34 | 507037 | LdCL_340017500 | amastin-like surface protein, putative | Gly6Ser |
| LdCL_34 | 1194752 | LdCL_340033400 | methyltransferase-like protein | Asp349del |
| LdCL_34 | 1547780 | LdCL_340044300 | Inositol-pentakisphosphate 2-kinase, putative | Val160Ala |
| LdCL_34 | 1777442 | LdCL_340051800 | sjogren s syndrome nuclear autoantigen 1, putative | Asp59Glu |
| LdCL_35 | 163307 | LdCL_350010100 | proteophosphoglycan ppg3, putative | Val2385Ile |
| LdCL_35 | 599964 | LdCL_350020200 | hypothetical protein | Cys10Tyr |
| LdCL_35 | 1412527 | LdCL_350042000 | hypothetical protein | Gly752Asp |
| LdCL_35 | 1716273 | LdCL_350050200 | hypothetical protein | Ile65Thr |
| LdCL_35 | 1716516 | LdCL_350050200 | hypothetical protein | Ala146Val |
| LdCL_35 | 1900240 | LdCL_350055800 | hypothetical protein | Leu120Ser |
| LdCL_36 | 958054 | LdCL_360030700 | hypothetical protein | Glu63Gly |
| LdCL_36 | 1300075 | LdCL_360040600 | oxidoreductase, putative | His269Arg |
| LdCL_36 | 1309078 | LdCL_360040900 | Fungal domain of unknown function | Arg64Leu |

| | | | | |
|---|---|---|---|---|
| | | | (DUF1712), putative | |
| LdCL_36 | 1354815 | LdCL_360042900 | CRAL/TRIO domain containing protein, putative | Thr468Ala |
| LdCL_36 | 1354866 | | | Pro451Ser |
| LdCL_36 | 1887706 | LdCL_360057700 | WD domain, G-beta repeat, putative | Gly416Glu |
| LdCL_36 | 2255530 | LdCL_360068100 | mkiaa0324 protein-like protein | Lys86Glu |

**Supplementary Table S2.**

Polymorphisms identified in the SL3 group were filtered by removing any SNPs in common with visceral *L. donovani* complex species and retained only if present in all three SL3 group isolates and in common with *L. major* alleles.
